# CoPR: Collective Pattern Recognition—a Framework for Microbial Community Activity Analysis

**DOI:** 10.1101/2024.06.30.601456

**Authors:** Rajith Vidanaarachchi, Sen Lin Tang, Saman Halgamuge

## Abstract

**Background:** Microbial community activities provide essential information on understanding bacterial communities. Unfortunately, they are generally not directly observable. We rely on longitudinal abundance profiles to get insight into microbial community activities. Often datasets do not have sufficient longitudinal sampling points to successfully apply our algorithms. Hence, in this paper, we are interested in analysing multiple datasets from similar environments to alleviate the aforementioned problem. Furthermore, we wish to see whether collective pattern recognition would enhance our understanding of microbial community activities.

**Results:** In this paper, we present CoPR, a framework for collective microbial longitudinal abundance data. Our visualisation shows that a single pattern for temporal abundance variation does not exist. However, it also indicates that even complete individuality does not exist. Consequently, our visualisation highlights the individuality and conformity in the temporal variation of abundance profiles of similar host environments. We also identify different characteristics in the TVAP (Temporal Variation of Abundance Profile) patterns with regards to cohesion and separation.

**Conclusions:** CoPR helps gain essential insights into the microbial communities and their heterogeneity through visualisation tools. This paper also highlights the choice between individuality and conformity in microbial community data analysis.

## 1 Background

We cannot generally observe microbial activity in the host environment and we rely on longitudinal observations of microbial abundance profile data to infer their activity. IMPARO [1], explored techniques for using mathematical models and optimisation to interpret the temporal variation of the abundance profile and infer interactions in the microbial communal activity.

In this paper, we continue our quest for answering the question of “What are they doing [in the microbial communities]?” [2]. In doing so, rather than quantifying the microbial behaviour as in the previous attempts, we look at the patterns in the temporal variation of the abundance profile (TVAP patterns). Visualising temporal variation in longitudinal experiments is a topic explored in recent words [3]. We can define a TVAP pattern as a particular pattern observable in the graph of abundance against time. It can be unique to a certain OTU or a specific host environment. We believe that comparing and contrasting TVAP patterns can infer insights into microbial community activity.

The inference of microbial activity through interaction inference algorithms is heavily reliant on the quality of available datasets. A useful dataset’s favourable qualities are high sampling frequency, consistent sampling frequency, and numerous sampled time points. Although many microbial abundance datasets are available [4], unfortunately, most of the available datasets do not feature these qualities.

For example, let us examine the Moving Pictures of the Human Microbiome study [5], which was analysed in this study was conducted over up to 15 months over 396 time points and provided time-series microbial abundance data of two individuals at four body sites [5]. The sampling frequency of this study is daily. We achieved high accuracy in inferring microbial activity in this dataset [1].

However, such datasets are not very commonplace. To illustrate the difficulty in collecting such a dataset, consider the scenario of a longitudinal study of the gut microbiome of a healthy individual. An individual available for daily stool analysis for six months would be the primary requirement of such a study. Furthermore, there will be barriers in terms of cost in studies that require specialist sample collection, such as coral microbiome collection requiring expert divers. On the other hand, some studies naturally have a shorter duration of interest, as in the menstrual vaginal microbiome—where the sampling needs to happen during the menstruation period [6].

Literature shows that many studies collected data parallelly from similar microbial communities–we consider microbial communities inhabiting the same type of host environments to be similar microbial communities. When a single dataset is not sufficient for inferring microbial communal activity, it has been shown that multiple datasets from similar communities could supplement the lack of data and provide more accurate inferences [7]. In this paper, we explore the idea of collective pattern recognition for enhancing our understanding of microbial community dynamics.

For example, one such dataset we explore in this paper comes from a study by [8]. They have sequenced the microbial communities in the guts of multiple premature infants in a neonatal Intensive Care Unit. We consider the premature infant gut as the type of host environment. Hence we use collective pattern recognition on microbial abundance profile datasets from all the infants in the study. With an average of 15 time-points per dataset, each dataset lacks sufficient data to successfully infer the microbial community dynamics. However, given that there are data from 58 infants, we can compare and contrast the different infants’ TVAP patterns and figure out common patterns of microbial behaviour—thus, this paper’s motivation is to combine datasets to recognise patterns collectively.

A second issue prevalent in analysing microbial community activities is the lack of independence from clinical or environmental factors. While the longitudinal abundance variation patterns reflect microbial activity (interactions), it is hard to discern the difference between the change in abundance due to internal—trophic or non-trophic—microbial activity and the change due to clinical factors and external influence. Even if the clinical data and a subset of environmental details are available, it is difficult to eliminate their factor into the abundance variation completely.

Using collective pattern recognition also allows discarding external factors up to an extent. Unless similar external factors affect all the host environments, collectively looking at the TVAP would identify patterns common to the host environment itself. If there are limited ways external factors affect the microbiome, collective pattern recognition allows identifying which host environments have been affected by the external factors.

Given the above reasons, we believe that collective pattern recognition will improve our understanding of microbial community activity.

### 1.1 Related Work

We explore related work under four main themes. Firstly, we look into the nature of the datasets available to gain insights into the requirements of working with similar data. Secondly, we look into microbial activity inference methods, as our end goal is to be facilitating the activity inference processes. Thirdly, we look into collective pattern recognition and clustering approaches, which plays a crucial role in our research. Fourthly, we look into the existing literature on individuality and conformity in microbial communities, under which we explore the ideas of microbial signatures, community state types, and precision medicine.

#### 1.1.1 Microbial Abundance Datasets

In this section, we will summarise some datasets where 16S rRNA sequencing has been used to collect time-series data from multiple similar host environments. *Premature Infants’ Gut Microbiome* [8] presents data from 58 neonatal infant gut microbial communities. This dataset was of interest as all the samples were collected while the infants were undergoing care at the neonatal intensive care units, which limited the gut microbiome’s interaction with the outside world, thereby limiting the external factors into the dynamics of the microbial community. They have collected 922 samples with an average of just over 15 per infant, up until 36 weeks of post-conception age, for all stool passings for each infant.

*Vaginal Microbiome of Reproductive-Age Women* [9] presents a dataset from 32 reproductive women’s vaginal microbial communities. They collected 937 samples over 16 weeks, with twice a week sampling frequency, averaging just over 29 samples per woman.

*Human Microbiome Related to Pregnancy* [10] have collected over 2500 samples from 49 pregnant women, pre– and post–delivery. They collected microbial community samples from the vagina, distal gut, saliva and tooth/gum just under 20 samples per site per woman. The collection frequency was weekly during gestation and monthly after the delivery.

*Neonatal Gut and Respiratory Microbiome* [11] have compiled another infant dataset from 82 infants. The data was collected over up to a year after birth with a weekly sampling frequency at the hospital and monthly thereafter. They have collected data from the gut (average of 13 per person) and respiratory tracts (nasal—an average of 12 and throat—an average of 6).

Availability of data of this nature makes our framework necessary and feasible.

#### 1.1.2 Microbial Community Activity Inference

Out of the many microbial community activity inference approaches, most depend on high-frequency datasets with a higher number of data points. IMPARO [1] uses an evolutionary algorithm with Lotka-Volterra equations to approximate microbial interaction parameters. MetaMIS [12] uses partial least square regression to estimate interaction parameters. Other methods include SparCC [13], which uses statistical methods, and the method by [7], which uses integrated data from multiple subjects in a Dynamic Bayesian network-based model.

The majority of the above methods use time-series data from a single host environment, and the results are significantly impacted by the availability of a large number of frequently sampled data points. Also, they do not utilise the availability of datasets with time-series samples of multiple similar host environments.

#### 1.1.3 Collective Pattern Recognition, Clustering, and Temporal Aligning Approaches

The idea of collective pattern recognition has been previously discussed in multiple works. Less so in the field of microbial interaction inference or related to microbial abundance data, but mainly in the area of gene expression analysis. As we can draw parallels between many biological data types, we will be looking at collective pattern recognition (including clustering and temporal aligning focused) work covering various applications.

1. [7] has explored collective pattern recognition for inferring microbial abundance patterns successfully. They align the temporal variation patterns coming from multiple host environments and define a typical pattern. They use this common pattern together with a dynamic Bayesian network to successfully predict microbial composition.
2. [14] reviewed clustering mechanisms to explore the response to external signals in time-series gene expression data. They also reviewed combining time-series data with other dynamic and static genomics data to better model gene expression patterns.
3. [15] used dataset alignment techniques and clustering to estimate unobserved data points in gene expression data. With datasets aligned by modelling them as piecewise polynomials, they had been able to achieve biologically meaningful results.
4. [16] also used time-series aligning techniques for temporal gene expression data. What makes their work interesting is that they present clustered data alignment, removing the assumption that all genes share the same alignment. Theirs is the first work to not treat gene expression data as homogeneous. Their inter-cluster independence in aligning temporal data provides more accurate alignments than earlier methods.
5. [17] used time warping algorithms when working with RNA and protein expression datasets. They show that time-warping clustering is superior to standard clustering using both interpolative and simple time-warping techniques. Their work is interesting in not assuming time-series biological data to be homogeneous in their temporal variation.
6. [18] also used time-warping techniques for alignment and template matching of time-series gene expression data. In their work, they have adapted dynamic time warping techniques from speech recognition research.
7. [19] introduce a statistical framework for co-expression networks. They use a kernel function to measure the similarity between subjects, which we identify as a collective pattern recognition technique. Their method, applied to time-series gene expression profiles of a group of subjects with respiratory virus exposure, produced early and accurate results.
8. [20] also analyses gene-expression data in their recent work. While using data from multiple subjects to model disease-relevant pathways, they also allow personalisation through a Gaussian process to identify differentially expressed genes. Their work claims to be more robust for identifying disease-relevant pathways in heterogeneous diseases.
9. [21] recently used time-series clustering techniques with lag penalisation for gene expression and protein phosphorylation datasets. They successfully identify clusters with distinct temporal patterns in both yeast osmotic stress response and axolotl limb regeneration studies. This study exemplifies that heterogeneous temporal variation behaviour is observed across various biological processes.
10. [22] used dynamic time warping techniques for comparative time-series transcriptome analysis in their recent work, TimeMeter. They were successful in characterising complicated temporal gene expression associations. They uncover exciting patterns in mouse digit restoration and axolotl blastema differentiation datasets.

#### 1.1.4 Individuality and Conformity

Individuality is identified as a microbial community behaviour, which is not uniform across communities of similar nature. Conformity is the opposite when similar OTU communities behave in a set predictable pattern. The individuality of microbial communities has been identified in the literature [23]. Especially in research on the gut microbiome, precision medicine has been proposed and successfully used in several studies [24–27]. Conforming behaviour has also been reported in the literature [28]. In the case of OTU communities, the concept of Community State Types [29] is an existing approach of explaining the balance of individuality and conformity in microbial communities.

Introduced by [29], [11], [10] and other studies report community state types (CSTs) in various microbial communities. The idea of CST is based on the composition of the constituent OTUs; as such, it is defined for a snapshot in time. [10] further notes that some communities show different state types, while some may show the same CST throughout the entire sampling period.

However, most microbial activity inference methods consider a single microbial community in their inference process, thus taking a highly individualistic approach. Some reasons for this could be the complexity of the external factors affecting the microbial community dynamics [1].

[7], however, uses a unified model obtained by aligning different microbial community TVAPs and assumes that a general microbial community of that particular type will take a particular pattern. [10] considers a vaginal community signature in their analysis of the vaginal microbiome.

Looking at the literature cited above, we can acknowledge that both approaches in considering individuality and conformity in the microbial community analysis process have their own merits.

### 1.2 Motivation and Contributions

It was interesting to note that collective pattern recognition goes hand-in-hand with individuality and conformity of microbial community dynamics. Considering the availability of studies where multiple temporal datasets of similar environments are available and the successful prior use of collective pattern recognition for biological data analysis, we further explored collective pattern recognition for microbial community dynamics analysis. We were motivated to use the collective pattern recognition techniques to shed light on individuality and conformity in microbial community dynamics and the heterogeneous nature of microbial abundance datasets.

We use unsupervised learning and visualisation techniques to analyse microbial abundance datasets and examine the TVAP patterns. We also talk about individuality and conformity, and heterogeneity of data in relation to microbial abundance datasets in our work.

In this paper, we present CoPR (Collective Pattern Recognition), a framework for analysing microbial community activities, which aims to address problems in lack of temporal abundance data and effects of external factors. CoPR primarily clusters OTU communities based on their TVAP patterns.

Ours is the first work to analyse the balance between individuality and conformity in microbial community activity patterns. Many existing works attempt to isolate a single pattern for temporal activity in microbial communities when working with multiple datasets. However, we consider microbial communities to be individualistic to a certain degree and look at multiple temporal activity patterns microbial communities can follow. Thus we believe our framework allows a more accurate analysis of microbial community activity.

CoPR also considers the heterogeneity of microbial datasets. We talk about microbial heterogeneity on multiple aspects and show that microbial abundance data, when treated as non-homogeneous, can uncover important details about the temporal community activity.

We present the analysis of multiple real-life datasets and simulated data. Our analysis identifies that the qualities of individuality and conformity in microbial communities are present across varying taxonomic resolutions, abundance levels and host environment types.

Key contributions summarised:

- Identifying correlations between different OTU TVAP pattern clusters.
- Framing the discourse around the balance between individuality and conformity, which we believe is essential to understanding microbial community activity.
- Exploration of heterogeneity of OTU TVAP patterns.
- Verification of the framework through the analysis of multiple real-life datasets (varying taxonomic levels, abundance levels, from different host environments, etc.) and simulated data.

## 2 Results

We processed several datasets—both real-life and simulated—through the pipeline and visualised patterns in the temporal variation of abundance profiles (TVAP patterns). Then we analysed the visualisations to illustrate how we can use them to gain insights into the microbial community dynamics. We present the results that illustrate some of the key arguments in this section. First, let us look at an example visualisation (Figure 1) to clearly understand the underlying meaning.

**Figure 1.**
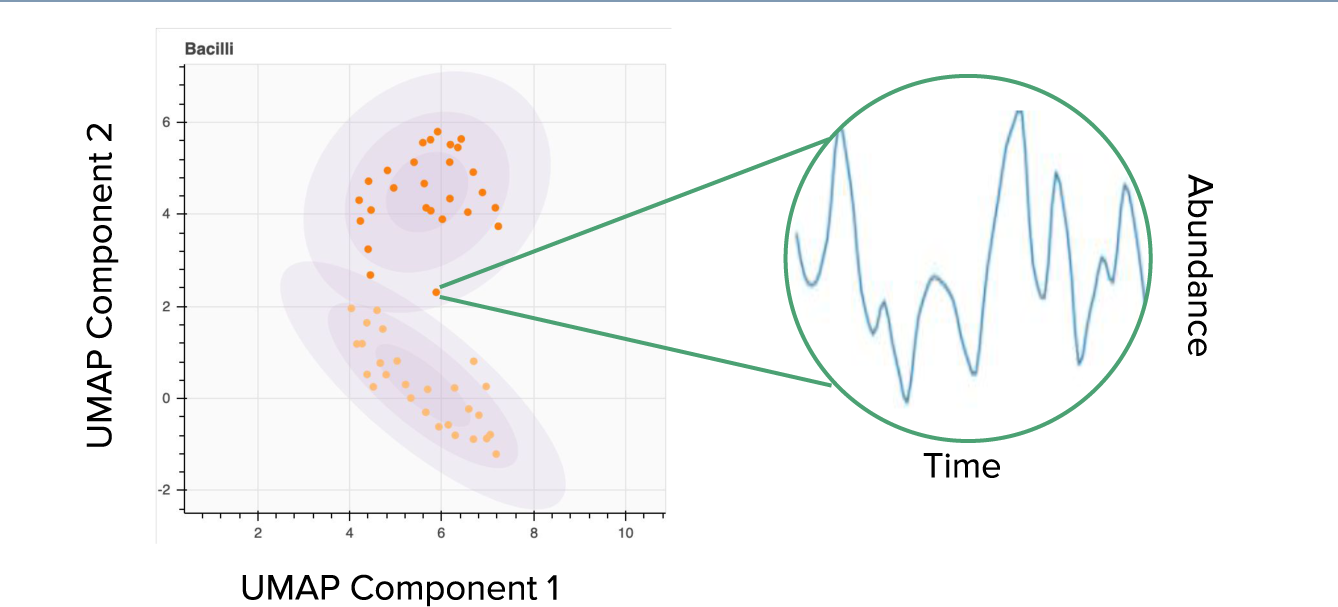
An example visualisation. In the cluster plots, each axis represents a dimensionally reduced component. Each dot in the cluster plot represents a specific environment’s temporal variation of the abundance profile (TVAP) of *Bacilli* (in this example). Host environments where *Bacilli* show similar TVAPs are clustered together, while host environments where *Bacilli* show distinct TVAPs are placed further away from each other. Trios of co-centred circles represent Gaussian Mixture Model clusters, where the varying opacity indicate the likelihood of a datapoint belonging to that cluster.

### 2.1 Non-Conformity Among the Communities of the Same OTU in Different Host Environments

We explore the temporal variation of abundance patterns of four OTUs—*Bacilli, Actinobacteria, Clostridia,* and *Gammaproteobacteria*—in the neonatal infant gut data set [8]. These particular OTUs were selected as they form the intersection of the ten highest abundant (averaged over time) OTUs in all the host environments (infants)—we shall call these the major OTUs (see Subsection 5.3).

The first observation we would like to draw attention to is how each OTU’s TVAP is separated into different clusters, as seen in Figure 2. This clustering indicates no conformity to a typical pattern observed for an OTU across different communities.

**Figure 2.**
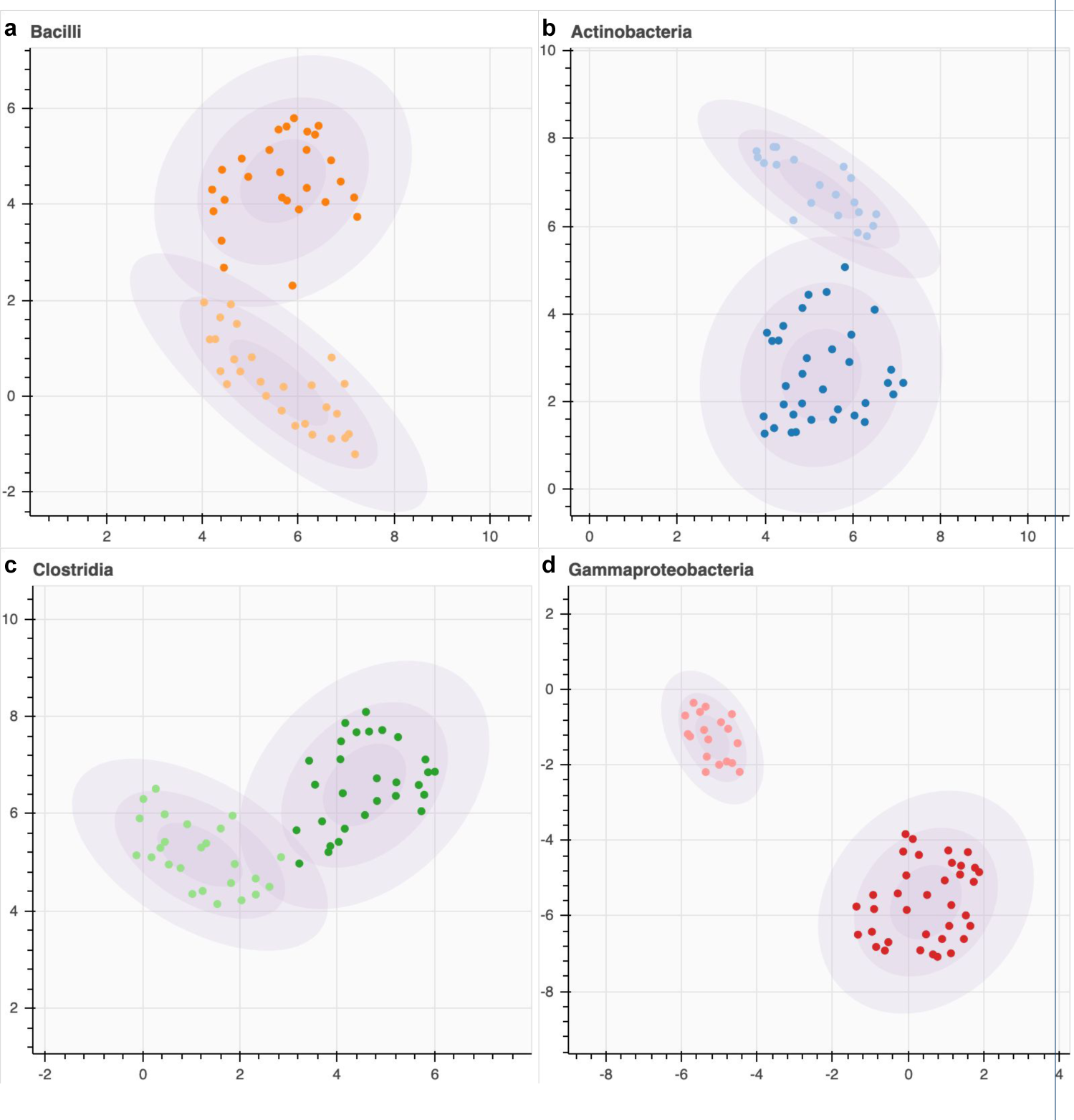
Clustering of TVAP of the major OTUs in the gut microbiome of preterm infants while in a neonatal ICU [8] is shown in the figure. Distances between TVAP were calculated using dynamic time warp (DTW) distance and visualised with UMAP (Uniform Manifold Approximation and Projection). Colours are local to each figure and represent clusters identified through Gaussian mixture models (GMM) clustering, where the highest silhouette score determined the number of clusters. All four major OTUs (a. *Bacilli*, b. *Actinobacteria*, c. *Clostridia*, and d. *Gammaproteobacteria*) show clear cluster separation. *Gammaproteobacteria* shows the most explicit separation. Although the optimal cluster number, according to silhouette score, is two, we observe sub-cluster separations in the dark red cluster. **The axes in these plots are: UMAP Component 1 (x) and UMAP Component 2 (y).**

This heterogeneous behaviour can again be identified in the TVAP curves seen in Figure 3.

**Figure 3.**
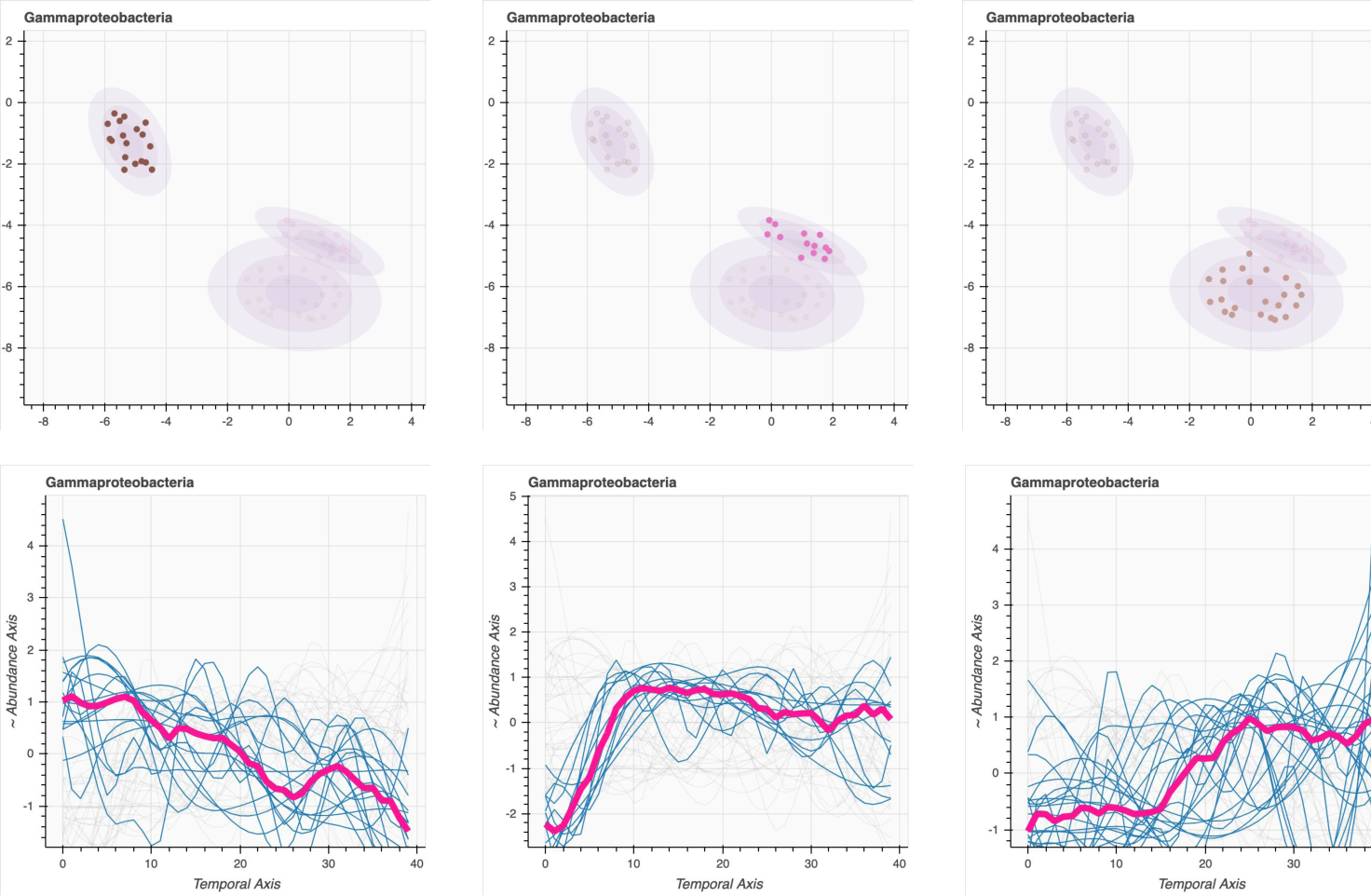
The figure shows distinctly identifiable TVAP patterns of *Gammaproteobacteria* in the [8] dataset. Each subfigure in the top row highlights a separable cluster of subjects, and corresponding subfigures in the bottom row show the TVAP of the highlighted subjects (blue lines). The thick pink line represents the median. The highlighted cluster in the third column shows more variation than the other two. It can be observed that the cluster from the first column encapsulates subjects in which the *Gammaproteobacteria*’s relative abundance drops from birth to discharge. In the second column, the relative abundance rises steeply soon after birth and maintains an abundance level thereafter. The third column is harder to classify clearly but shows a general trend of rising and falling while favouring a final rise. **The axes in the top row plots are: UMAP Component 1 (x) and UMAP Component 2 (y).**

### 2.2 Conformity Among the Communities of the Same OTU

Secondly, we observe that the different communities neither subscribe to a typical behaviour nor completely sporadic. Let us consider the clusters shown in Figure 2. Especially the clusters of *Gammaproteobacteria* are tightly knit together. Although we have clustered the *Gammaproteobacteria* communities into two clusters according to the silhouette value (see Secion 5), we observe a distinctly identifiable subcluster within one of the main clusters. In Figure 3, we have increased the cluster numbers to closely observe how the temporal variation patterns differed. It indicates that although the primary separation is based on falling–rising behaviours, the sub-clusters differ on when the rise happens. The communities belonging to the smaller of the three clusters all show an initial rise in *Gammaproteobacteria* abundance. This cluster, along with the cluster in the third column of Figure 3, where a general rising behaviour is observed, formed the larger cluster of Figure 2 (indicated in deep red). Interestingly this conformance to a specific behaviour happens in certain subsets of communities.

Observing the TVAP plots in Figure 3, we also note that the deviancy from the cluster’s median is different for each cluster. Observation of this heterogeneous behaviour is also of interest.

However, as a summary, we can say that there are three distinctly identifiable TVAP patterns amongst *Gammaproteobacteria* communities in the infant gut. The first is a gradual reduction of relative abundance; the second is an initial rise in relative abundance and subsequent maintenance. The third can be categorised as a late rise in relative abundance. In our original clustering of Figure 2, the two clusters represented a rise and a fall in relative abundance. In the finer clustering of Figure 3, the rise was characterised into two different rising patterns. Although this behaviour is demonstrated in the *Gammaproteobacteria* communities, sub-cluster separation can be identified in other communities as well. We can discern the existence of possibly separable sub-clusters by observing silhouette scores.

### 2.3 Agreement of Clusters of Different OTUs

Next, we investigated the connection between the clusters of different OTUs. To quantify this, we looked at the overlap coefficient (see Section 5) of cluster membership. The cluster membership distribution calculated for the same dataset can be seen in Figure 4. Firstly, it gives us a quantitative sense of the overlaps, and secondly, it provides us with information that is hard to discern visually.

**Figure 4.**
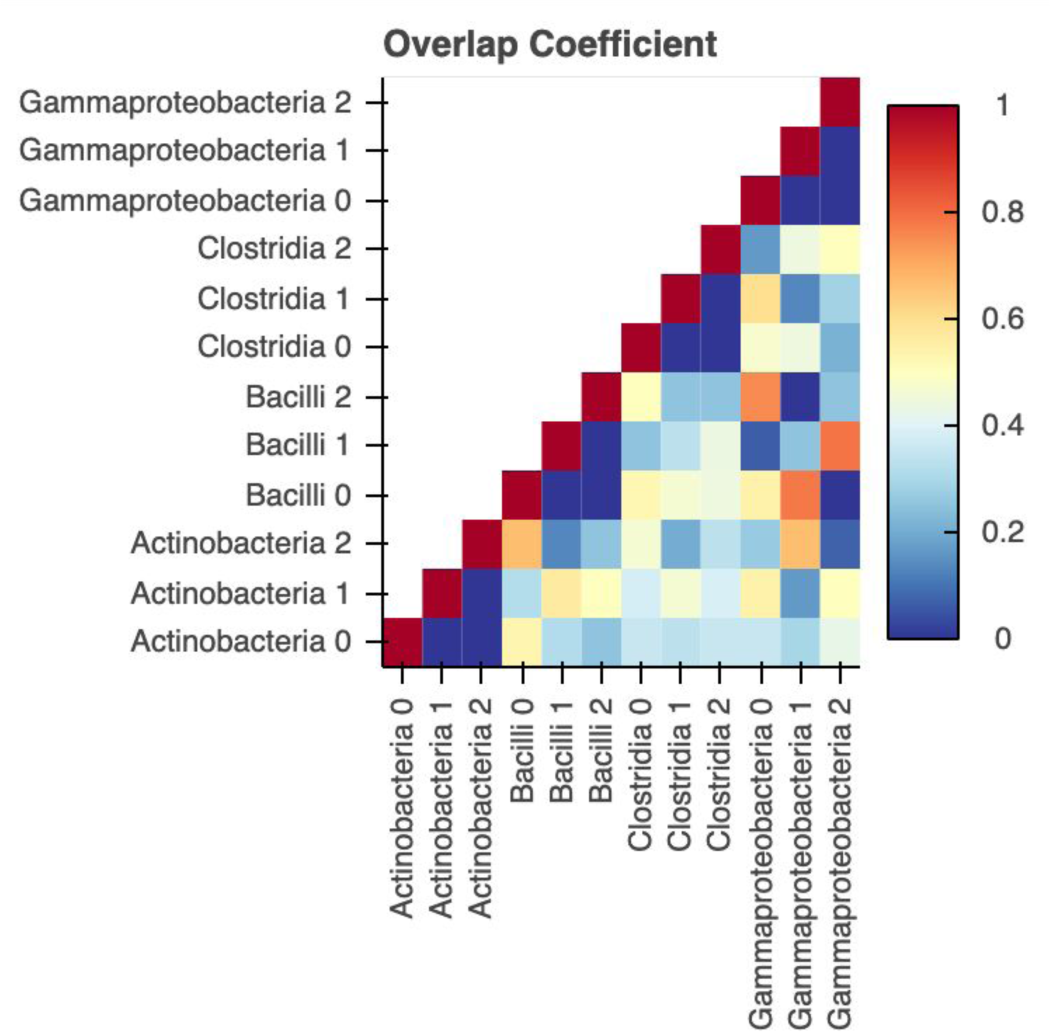
This figure shows the distribution of the overlap coefficient between the different clusters in the analysis of the [8] dataset. *Gammaproteobacteria*, *Clostridia*, *Bacilli*, and *Actinobacteria* were clustered into three. Highest overlaps are observed in *Gammaproteobacteria* and *Bacilli* clusters.

From Figure 4, we identified the overlaps of *Gammaproteobacteria* and *Bacilli*. This association between the three clusters were statistically significant, *χ*^2^ = (4*, N* = 54) = 43.6075, *p* = 7.40076*E −* 9 (Fisher’s exact test = 1*E −* 8) (*<* 0.01).

Furthermore, the relationship between the two main clusters of Gammaproteobacteria and Bacilli were statistically significant, *χ*^2^ = (1*, N* = 54) = 10.7628, *p* = 0.001036(*<* 0.01). We further explored the corresponding behaviour in Figure 5. In the left half of the figure, we observe that *Gammaproteobacteria*’s TVAP rises initially and maintains that level, while *Bacilli*’s corresponding TVAP falls and maintains very low. Interestingly while this exchange of prominence in the bacterial community is clear, we cannot find a clear explanation for the corresponding behaviour in the other two clusters from the median TVAP patterns.

**Figure 5.**
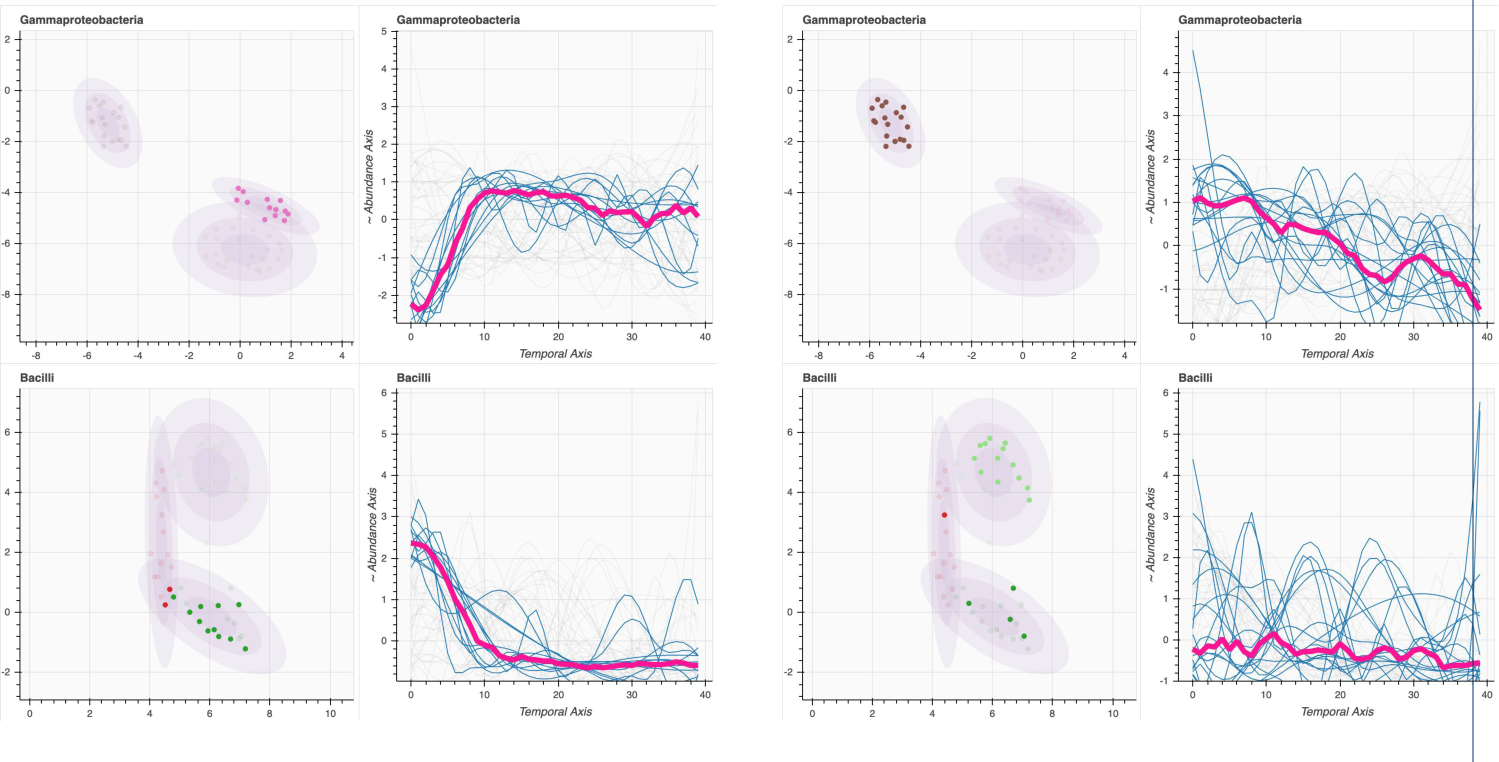
The figure shows the corresponding behaviour of *Gammaproteobacteria* (top row) clusters with *Bacilli* (bottom row) clusters. Subfigures show TVAP clustering and the TVAP curves corresponding to the highlighted host environments (infants), with the median curve for the subset of highlighted host environments shown as the thick pink line. Clusters on the left show that the initial rise in Gammaproteobacteria corresponds with an initial fall in Bacilli (The overlap coefficient for these two clusters are 71% with a Jaccard Index of 48%). Clusters on the right show that decreasing relative abundance of Gammaproteobacteria corresponds with the rising and falling behaviour of Bacilli. Observations seen in the clusters on the left are in line with the observations of [8]. Still, while they generalise this observation for the whole community, we highlight that this is only one of the three distinctly identifiable trends observable in this dataset. **The axes in the cluster plots are: UMAP Component 1 (x) and UMAP Component 2 (y).**

### 2.4 Clusters and External Factors

We were interested in investigating whether the clustering of host environments had a connection to the external factors. Some datasets include clinical factors and other environmental variables, which could broaden the value of our analysis. For example, in our analysis of the neonatal gut microbial dataset [8], a connection between the infants’ delivery method seemed to correlate to the clustering of *Bacilli* and *Gammaproteobacteria* TVAPs. However, a chi-square test of independence showed no significant association between the TVAP clustering and delivery method. *χ*^2^ = (1*, N* = 54) = 0.8105, *p >* 0.1

### 2.5 Common Themes in Multiple Real Life Datasets

We explore a second dataset, which consists of gut, nasal and throat microbiomes of infants [11]. The primary aim of this exploration is to observe common themes which can be identified in real-life datasets.

In Figure 6, we note that six major OTUs are identified in the [11] gut dataset. Among the six are the four identified in the [8] study’s analysis. The separation of clusters also shows similarity, with *Gammaproteobacteria* communities arguably separating well, although not as well as in the previous study. *Actinobacteria* is also notably separated into three different clusters. However, overall, in all the OTU *clusterings*, we can identify many clear subclusters. We also observe connected clusters in *Bacteroidia* and *Coriobacteriia* communities.

**Figure 6.**
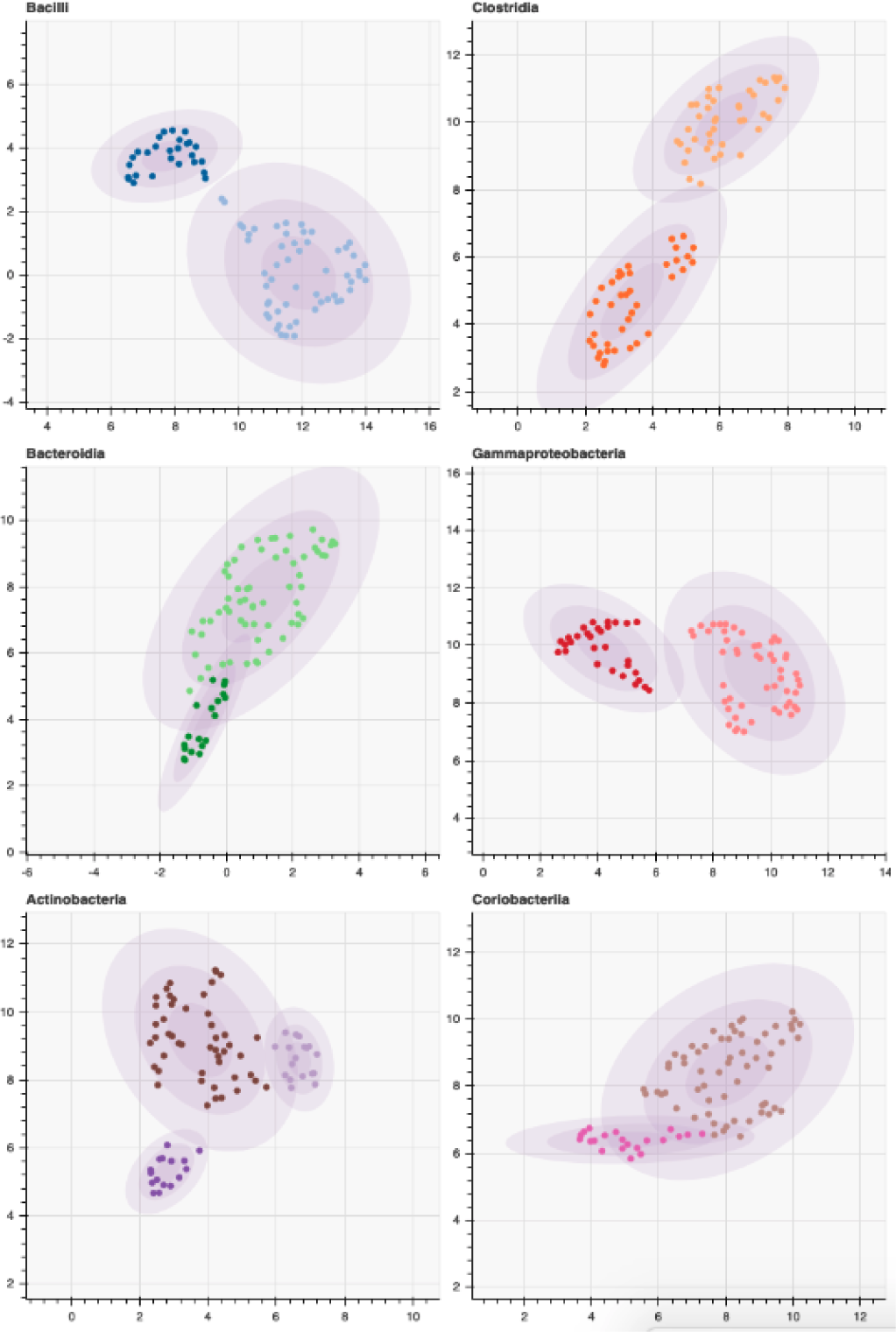
Clustering of TVAP of the major OTUs in the gut microbiome of infants from [11] study are shown in the figure. Distances between TVAP were calculated using DTW, and and visualised with UMAP. Colours are local to each figure and represent clusters identified through GMM clustering, where the highest silhouette score determined the number of clusters. All four major OTUs (*Bacilli*, *Gammaproteobacteria*, *Clostridia*, and *Actinobacteria*) found in the [8] dataset are also found to be major OTUs in this dataset, with the addition of *Corriobacterlia* and *Bacteroidia*. Similar to the [8] study, *Gammaproteobacteria*, *Bacilli*, and *Clostridia* show clear separation into two clusters. **The axes in these plots are: UMAP Component 1 (x) and UMAP Component 2 (y).**

In Figure 7, we explore the TVAP patterns in the nasal microbiome from the [11] study. We observe five major OTUs identified, of which *Clostridia*’s separation and *Betaproteobacteria*’s separation are clearly identifiable. Most noteworthy out of all is *Bacilli*’s TVAP patterns, which show a set of triangularly interconnected clusters. This cluster layout is an excellent example of the gradual behavioural change in the communities where a balance between individuality and conformity may exist. *Bacteriodia* and *Actinobacteria* show connected clusters as well.

**Figure 7.**
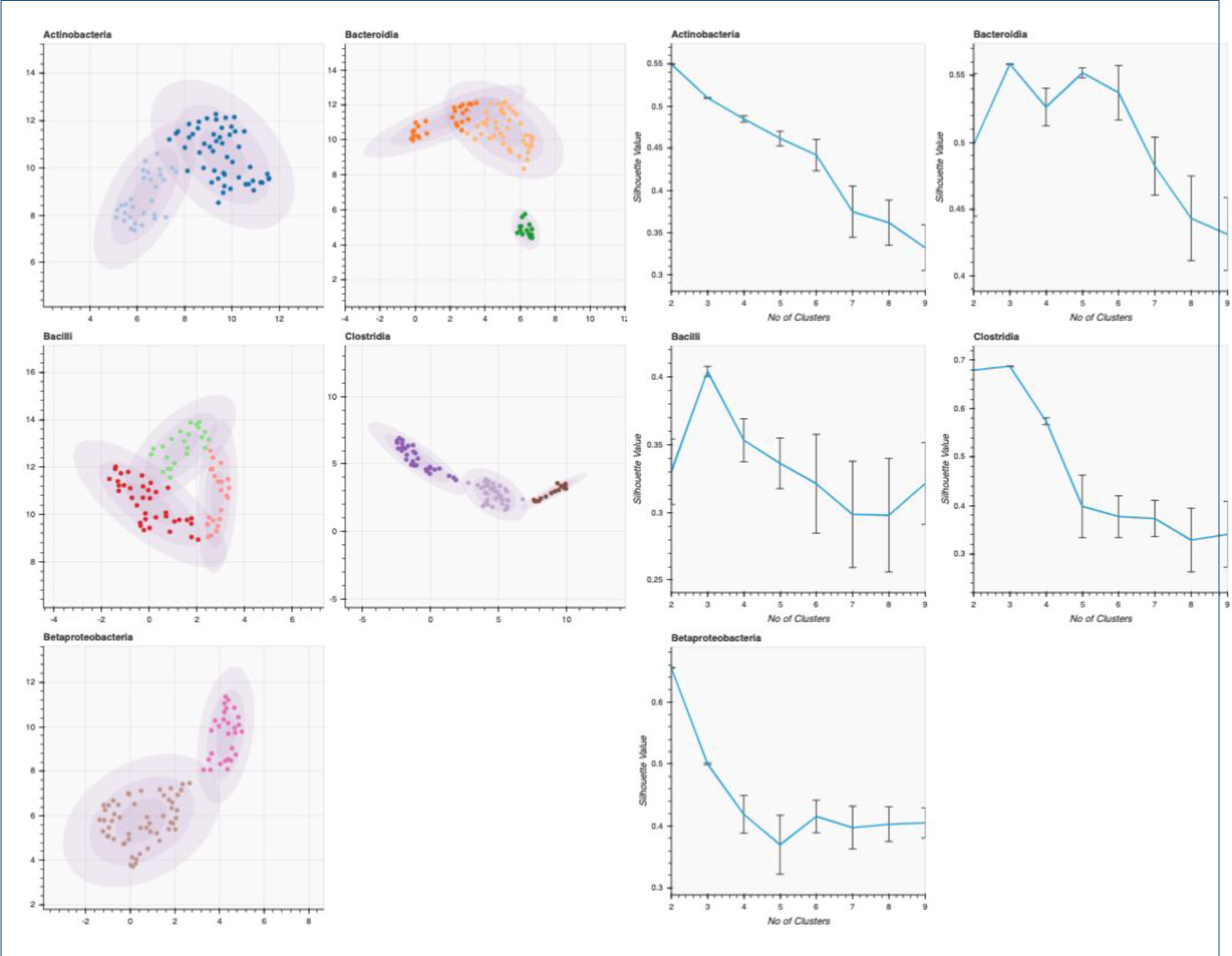
Clustering of TVAP of the major OTUs in the nasal microbiome of infants from [11] are shown in the left-hand figure. Distances between TVAP were calculated using DTW and visualised with UMAP. Silhouette scores for different cluster numbers for each of the major OTUs are plotted in the right-hand figure. Colours are local to each sub-figure in the left-hand figure and represent clusters identified through GMM clustering. The number of clusters was determined by the highest silhouette scores, as shown in the right-hand figure. Three major OTUs (*Bacilli*, *Clostridia*, and *Actinobacteria*) found in the [8] dataset are also found to be major OTUs in this dataset. The absence of *Gammaproteobacteria* could be explained by the aerobic environment of the nasal cavity. While some OTUs have silhouette scores indicative of a superior number of clusters, others have closely competing cluster numbers. **The axes in the cluster plots are: UMAP Component 1 (x) and UMAP Component 2 (y).**

In Figure 8, we explore the TVAP patterns of the throat microbiome of [11], where six major OTUs have been identified. We observe clearly disjoint clusters in *Fusobacteria* and *Betaproteobacteria*. Throat microbiome is the second type of community where *Betaproteobacteria* shows good separation, and the third, when we consider *Proteobacteria* as a whole (Figures 6, 7 and 8). The cluster shapes and directions again prove interesting, with *Clostridia* showing a tree-like structure. This structure suggests that each cluster shows gradual changes in three different directions, deviating from a central pattern

**Figure 8.**
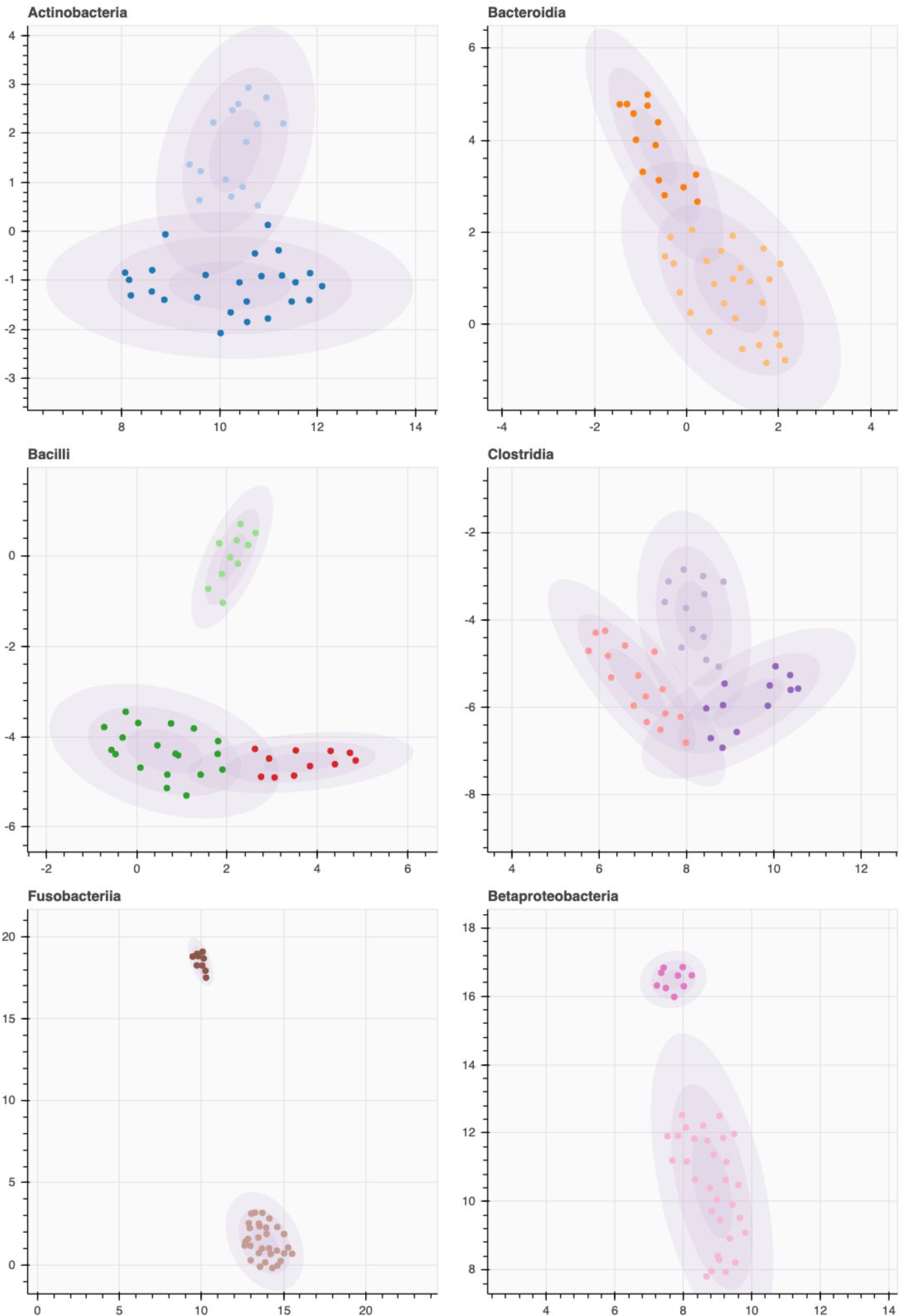
Clustering of TVAP of the major OTUs in the throat microbiome of infants from [11] study are shown in the figure. Distances between TVAP were calculated using DTW, and visualised using UMAP. Colours are local to each figure and represent clusters identified through GMM clustering, where the highest silhouette score determined the number of clusters. Three major OTUs (*Bacilli*, *Clostridia*, and *Actinobacteria*) found in the [8] dataset are also found to be major OTUs in this dataset. The absence of *Gammaproteobacteria* could again be explained by the aerobic environment of the throat cavity. While *Bacilli*, *Betaproteobacteria* and *Fusobacteria* show clear separation, other OTUs show gradual changes in TVAP. **The axes in the cluster plots are: UMAP Component 1 (x) and UMAP Component 2 (y).**

### 2.6 Differentiating Disjoint Clusters and Connected Clusters

A secondary observation we can make from Figures 2 and 3 is that in some OTUs, the clusters are distinct and disjoint, while in other OTUs, the clusters are connected. The same behaviour is highlighted in Figure 9. Here we can identify another peculiar behaviour of TVAP patterns: Disjoint clusters represent TVAP patterns which correspond to a set behaviour, as opposed to connected clusters, we can see a gradual change of behaviour in TVAP.

**Figure 9.**
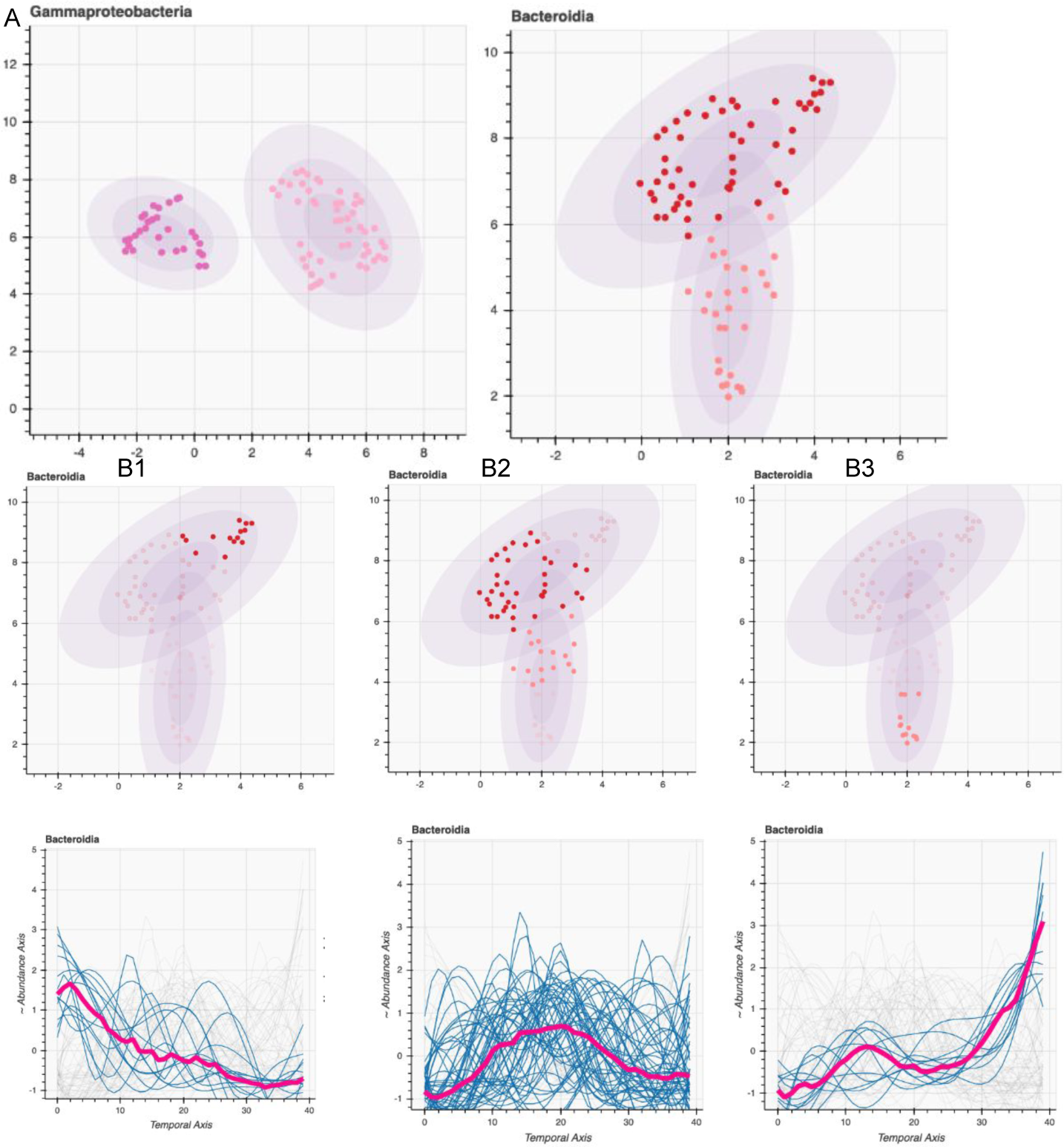
The figure shows clustering of the major OTUs in infants’ gut microbiome from the [11] study. Distances between TVAP were calculated using DTW and visualised using UMAP. Colours are local to each figure and represent clusters identified through Gaussian Mixture Models (GMM) clustering, where the highest silhouette score determined the number of clusters. The TVAP patterns of *Gammaproteobacteria* show separated two main clusters and subclusters within. However, the TVAP of *Bacteriodia* shows gradual change across the clusters, as shown in the subfigures B1 - B3. **The axes in the cluster plots are: UMAP Component 1 (x) and UMAP Component 2 (y).**

### 2.7 Analysis Across Taxonomic Resolutions

In Figure 10, starting from the phylum (L2) taxonomic level, moving up to genus (L6) taxonomic level, we have demonstrated that the traits of individuality and conformity are present at varying taxonomic levels—in addition to the observations at the class (L3) level. Also, we observe the TVAP pattern clusters change as we traverse through the taxonomic hierarchy. Also noteworthy is the conserved structure of the taxonomic hierarchy of Phylum *Firmicutes*, Class *Bacilli*, Order *Lactobacillales*, Family *Streptococcaceae*, and Genus *Streptococcus* which is featured in Figure 11. Also, at each taxonomic level, there are both disjoint and connected clusters present.

**Figure 10.**
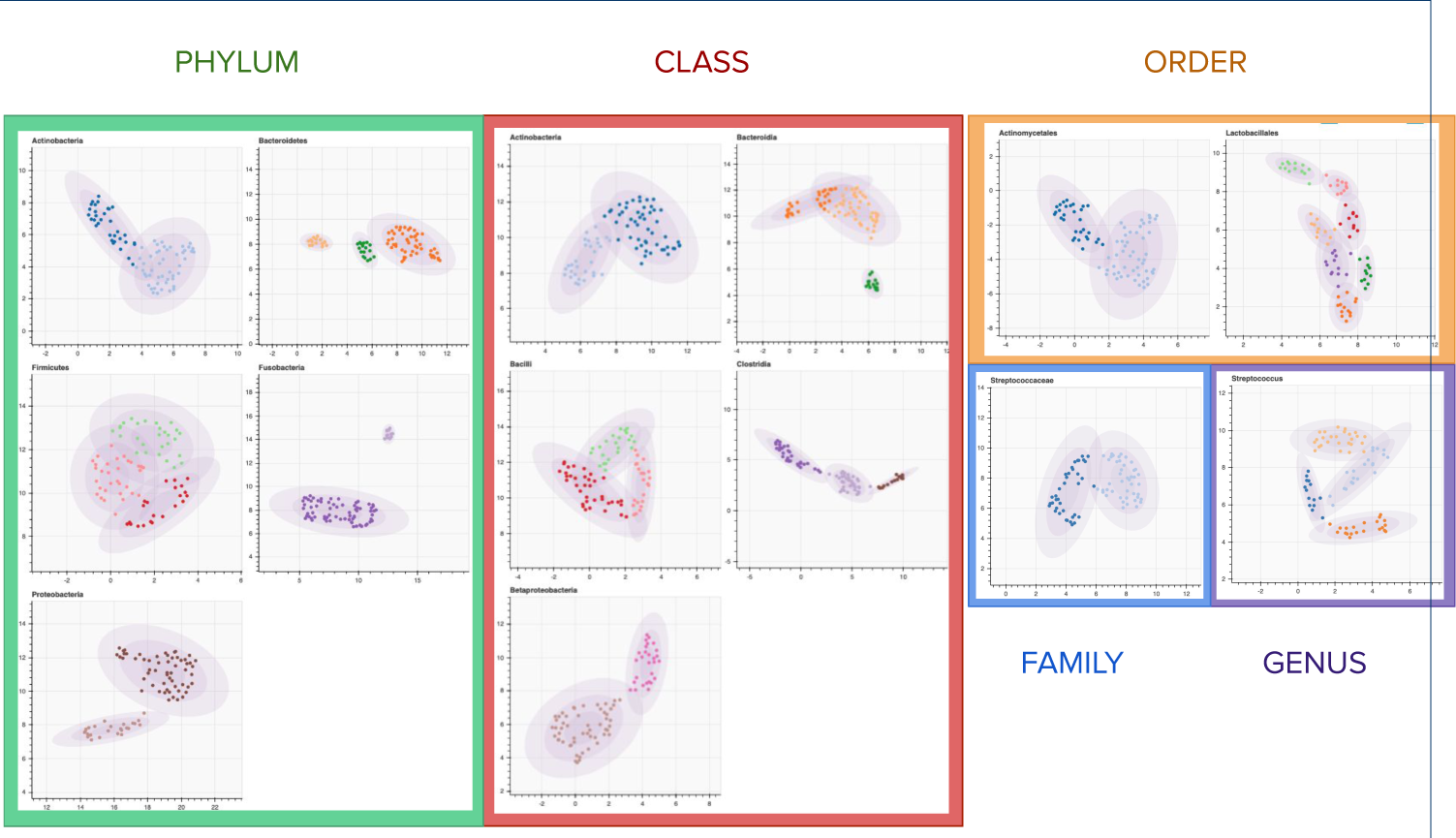
This figure shows the TVAP clustering of the infants’ nasal microbiome from [11] at various taxonomic levels. Colours are local to each figure and represent clusters identified through GMM clustering, where the highest silhouette score determined the number of clusters. The number of major OTUs reduces with the increased taxonomic resolution. However, at each resolution, we observe separated clusters. **The axes in the cluster plots are: UMAP Component 1 (x) and UMAP Component 2 (y).**

**Figure 11.**
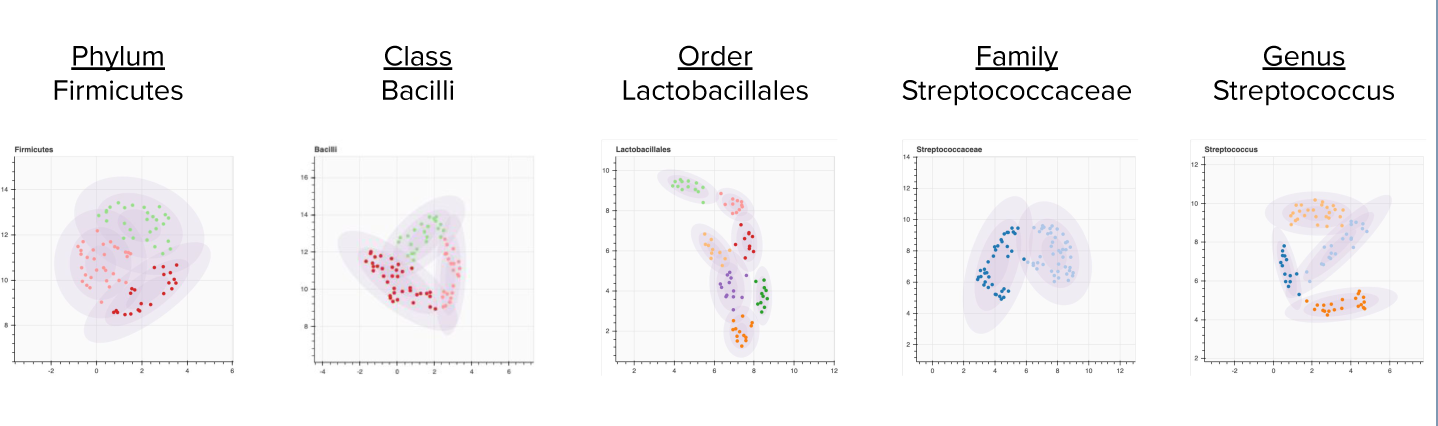
A selection along a branch of the taxonomic tree from Figure 10. We note change of the cluster structures across varying taxonomic levels. **The axes in the cluster plots are: UMAP Component 1 (x) and UMAP Component 2 (y).**

### 2.8 Major OTUs and Secondary OTUs

We mainly focused on the major OTUs, which were defined as the most abundant OTUs common to all the host environments. However, we are also interested in observing the nature of secondary OTUs’ TVAP patterns. Figure 12 shows a data set from the throat microbiome again, but we have chosen the second most abundant taxa instead of the major ones. We notice that even the secondary OTUs show attributes we discussed above. We especially take note in *TM7-3* and *Flavobacteria* which show clearly disjoint clusters. Although non-major OTUs are discarded in some studies from the analysis [12], we suggest that CoPR can successfully give meaningful visualisations for those.

**Figure 12.**
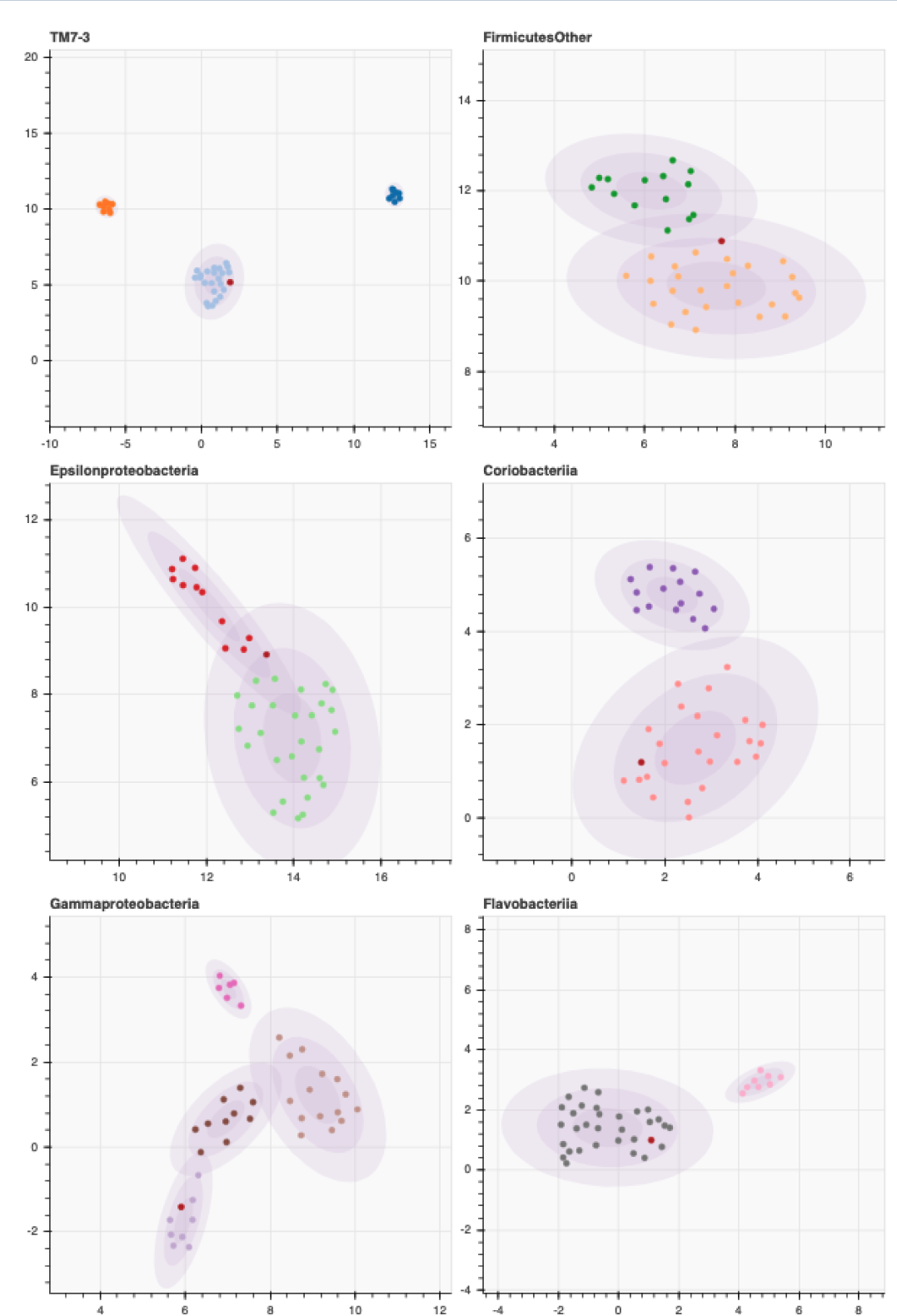
TVAP clustering of the secondary OTUs. Again, we are visualising the infant throat microbiome from [11] at Family (L3) taxonomic level, but with non-major OTUs; Colours are local to each figure and represent clusters identified through GMM clustering, where the highest silhouette score determined the number of clusters. Secondary OTUs also show similar behavioural patterns to those of major OTUs. **The axes in the cluster plots are: UMAP Component 1 (x) and UMAP Component 2 (y).**

### 2.9 Silhouette Scores and the Number of Clusters

In using any kind of clustering, deciding the optimal number of clusters is essential. We used the silhouette scores for this purpose. In Figure 7, we show the silhouette scores next to the cluster visualisations. We observe that silhouette score graphs can also bring invaluable information about the microbial community activity analysis. As an example, we will look at *Clostridia* and *Betaproteobacteria*. Each of these has clearly separated clusters, and clearly prominent peaks of silhouette scores at their respective optimal cluster numbers show that the clustering is robust. We also observe the standard deviation (error) bands at the respective optimal cluster numbers are small for these OTUs.

Another example is *Bacteroidia*, whose silhouette score peaks at three, but it also indicates that smaller cluster numbers can also provide “good enough” silhouette values. However, the standard deviation bands reconfirm that the clustering into three is the most robust and consistent option, regardless of the initial points selected for the clustering.

### 2.10 Simulated Data

After examining two real-life datasets, we look at a simulated dataset. The simulated dataset is created to approximate a known grouping with the clustering. Although ground-truth cluster labels are impossible to find in practice, we carry out the simulation to test the limits of CoPR in uncovering the known truths. In creating the simulated data, we faced several challenges. Foremost, it is currently impossible to create a dataset where the shape or the pattern of the TVAP is directly linked to the interaction parameters while preserving randomness. Hence we created multiple stencils for TVAP patterns and approximated them with known functions with randomness in parameters and noise. The experiment was designed to achieve the original stencil patterns as the median TVAP of each cluster.

The objective of the simulation was to approximate a typical longitudinal abundance dataset. The simulation has 100 subjects in total, with 20 OTUs in the microbial community. This number was much lower than what one would find in a typical microbial community. However, as most OTUs in a typical community are rare OTUs, we were satisfied with the simulation data generation process’s lower number. From the 20 OTUs, four were considered to be major OTUs, while the other 16 were of secondary abundance. Approximately a major OTU’s abundance was ten times that of a minor OTU. Out of the major OTUs, two—OTU-A and OTU-B—had two TVAP patterns each (Figure 13). Grouping of TVAP patterns in OTU-A and OTU-B corresponded to each other, except for randomly generated outliers. These outliers are highlighted in the TVAP cluster subplot of OTU-B corresponding to rising behaviour (Figure 13). OTU-C and OTU-D had three distinct behavioural patterns, which had no connection to each other or to that of OTU-A or OTU-B.

**Figure 13.**
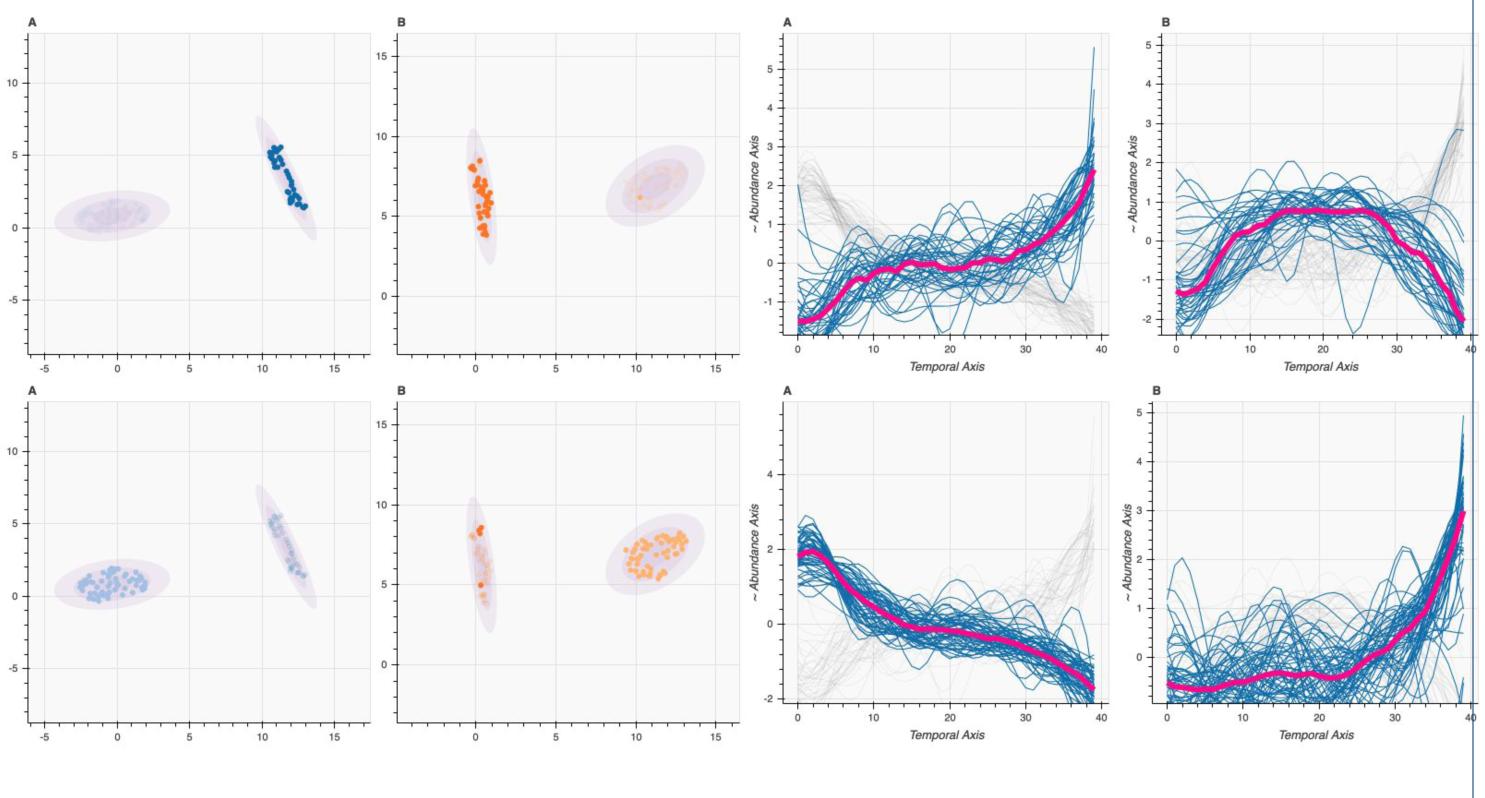
This figure shows the visualisation of the simulated OTUs A and B, with two corresponding clusters selected. The left-most half contains the cluster placement in the reduced dimension scatter plot, and the right-most half shows the TVAP of the selected clusters and the median for each cluster shown in thick pink. **The axes in the cluster plots are: UMAP Component 1 (x) and UMAP Component 2 (y).**

## 3 Discussion

In this section, we further discuss the concepts of individuality and conformity, analyse the visualisations obtained by CoPR, and discuss the significance of the key findings.

### 3.1 Individuality versus Conformity

Individuality is a quality, trait or behaviour that separates something from the rest. In contrast, conformity is the opposite, where something’s qualities, traits and behaviours are according to a set norm or standard and non-deviating from the rest. This concept has been applied in social science to describe human populations [30, 31]. These concepts have emerged briefly in the literature of ecology [32] and made a comeback only recently in the field of microbiology [33]. Inspired by these ideas, we would like to define the individuality and conformity of OTU activity in the scope of this work.

Let us define individuality as the tendency of the same OTU in similar environments to show different temporal variations in their abundance profiles. Conversely, let us define conformity as the tendency of the same OTU in similar environments to show similar variations in their abundance profiles. In the scope of this work, we define the traits of individuality and conformity to co-exist. We propose a fuzzy interpretation of the concepts, where each OTU community show a membership towards individuality and conformity.”

Individuality and conformity are not phenomena limited to the microbial world. We observe this in human society, animal and plant kingdoms and many other natural and human-made systems. In most of these systems, we encounter generalisations to be helpful to an extent. However, generalisations have to be considered, coupled with the correct assumptions of circumstance. While generalisations are helpful, generalisation beyond reasonable assumptions is not. As an example, take plant care. While it can be assumed that plants of the same species need similar care in most cases, there may be individual plants that require a different kind of approach. We hypothesise that reasoning similar to this is valid for the microbial world as well. When we develop a generalised model for microbial community dynamics, it is essential that we are aware of the singularities of each community. Our visualisations provide qualitative insight into this balance.

Identifying these common tendencies or conformities will help us build better models to simulate microbial communities in general. They will help us understand better the links between different OTU communities and different types of host environments. After identifying the conformities, we can also identify the individualities for further analysis. Together with the conformities, the individualities give us specialised information about the specific issues related to a single OTU community. Information on both levels will assist us in obtaining a more practical idea of the microbial community dynamics.

Firstly, we simply do not see everything gathered in a single cluster in the reduced dimension scatter plots. If that were the case, it would mean every community of the same OTU behaves in the same way. The OTUs being scattered around signify that there is no set norm for OTU communities’ activity in a specific host environment— it would be incorrect to assume that, for example, TVAP of *Clostridia* will always show a particular tendency in a human gut environment.

Conversely, we do observe clusters rather than completely scattered points in the reduced dimension scatter plots. This observation means that subsets of OTU communities do show similar behaviour. When an OTU shows multiple prominent clusters, we consider that there may be multiple likely ways for this particular OTU communities to behave in this particular host environment. Hence, the existence of clusters is a degree of conformity we observe in the microbial communities.

We further hypothesise a connection between individualistic traits and external factors. Each host environment has specific environmental, clinical or other external factors. These external factors certainly affect the microbiome and its dynamics. The concepts of individuality and conformity may very well indicate the communities’ reaction to their environment. More individuality than conformity in a particular microbial community’s behaviour may indicate the community’s sensitivity to the external environment. Although CoPR visualisation does not grant knowledge qualified to make a statement about the causality or correlation of the clinical factors to the microbial dynamics, we can conclude that these correlations can be identified. We summarise the observations from the results, together with the identified individualities and conformities in Table 1. In this table, we detail how specific observations can indicate individuality, and others, conformity. We would like to draw attention to concurrent observations that suggest both individualist traits and conformist traits.

**Table 1:**
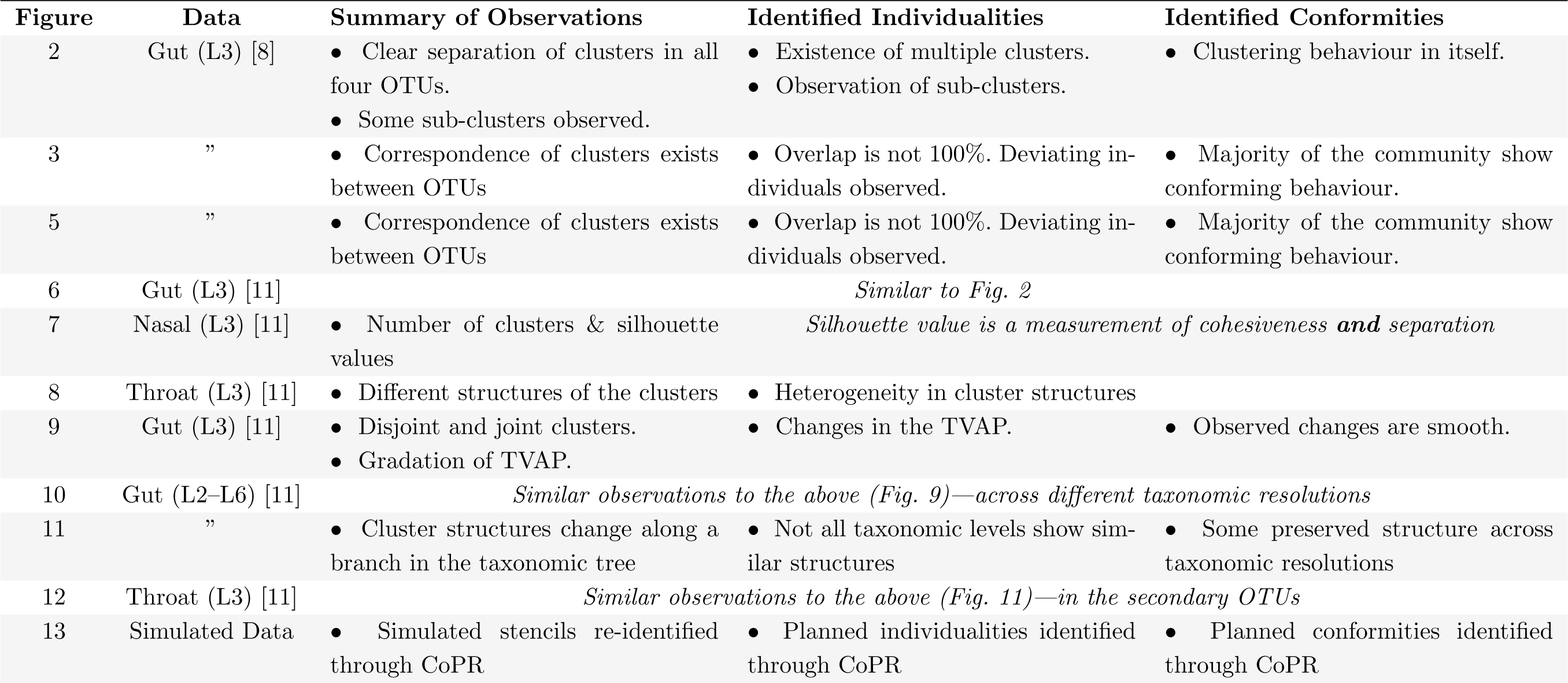
Summary of the Figures.

| Figure | Data | Summary of Observations | Identified Individualities | Identified Conformities |
| --- | --- | --- | --- | --- |
| 2 | Gut (L3) [8] | • Clear separation of clusters in all four OTUs.<br>• Some sub-clusters observed. | • Existence of multiple clusters.<br>• Observation of sub-clusters. | • Clustering behaviour in itself. |
| 3 | ” | • Correspondence of clusters exists between OTUs | • Overlap is not 100%. Deviating individuals observed. | • Majority of the community show conforming behaviour. |
| 5 | ” | • Correspondence of clusters exists between OTUs | • Overlap is not 100%. Deviating individuals observed. | • Majority of the community show conforming behaviour. |
| 6 | Gut (L3) [11] | *Similar to Fig. 2* |  |  |
| 7 | Nasal (L3) [11] | • Number of clusters & silhouette values | *Silhouette value is a measurement of cohesiveness **and** separation* |  |
| 8 | Throat (L3) [11] | • Different structures of the clusters | • Heterogeneity in cluster structures |  |
| 9 | Gut (L3) [11] | • Disjoint and joint clusters.<br>• Gradation of TVAP. | • Changes in the TVAP. | • Observed changes are smooth. |
| 10 | Gut (L2–L6) [11] | *Similar observations to the above (Fig. 9)—across different taxonomic resolutions* |  |  |
| 11 | ” | • Cluster structures change along a branch in the taxonomic tree | • Not all taxonomic levels show similar structures | • Some preserved structure across taxonomic resolutions |
| 12 | Throat (L3) [11] | *Similar observations to the above (Fig. 11)—in the secondary OTUs* |  |  |
| 13 | Simulated Data | • Simulated stencils re-identified through CoPR | • Planned individualities identified through CoPR | • Planned conformities identified through CoPR |

### 3.2 Visualisation

In this subsection, we will discuss the visualisations available from the analysis.

#### 3.2.1 GMM Clusters

Dimension reduction has been successfully employed to visualise high-dimensional datasets [34]. The main visualisation of our analysis is the Gaussian Mixed Model (GMM) Clustering. GMM is capable of identifying non-circular clusters as well. The EM algorithm that determines the cluster membership results in clusters of TVAP suitable for further interpretation in the context of microbiology. Each OTU is plotted separately, with each host environment represented by a dot plotted in the UMAP reduced dimensions. The UMAP dimension reduction is based on the Dynamic Time Warp distance between the TVAP curves. The plots also indicate cluster membership at three different membership probabilities. This visualisation primarily gives us information on how many distinct TVAP patterns could be identified for the specific OTU and how separated they are from each other. Secondarily we can identify cluster placement and shapes, which can provide us with information about the microbial dynamics.

#### 3.2.2 Median Plot

The median plot of the TVAPs is straightforward in terms of mathematics associated but is very useful in comparing and contrasting the temporal variation across different clusters. The median line also helps us visualise the deviation of the member OTU’s patterns from the typical pattern. We may pass on the median TVAP lines from all the OTUs to a microbial interaction inference algorithm to obtain interaction parameters.

#### 3.2.3 Silhouette Index

Silhouette Index, plotted against the number of clusters, provides insight into the nature of the clusters. The silhouette index provides a quantitative measure of the clusters’ cohesion and separation, reflecting the microbial communities’ individuality and conformity. This quantification complements the qualitative idea we gather from the cluster plots. The error bars in the silhouette index plots indicate how consistent the clustering is.

#### 3.2.4 Jaccard and Overlap Indices

Jaccard and Overlap indices are suitable metrics to confirm the visual from the cluster plots as they are quantitative measures of the agreement between the clusters of different OTUs. These plots complement the cluster plots.

### 3.3 Assumptions Involved

In any method involving microbial dynamics, there are several assumptions involved. These assumptions are due to the unobservable nature of the microbial dynamics and the poor understanding of various internal and external factors that affect the microbial communities. We also take the liberty to involve some assumptions in our application pipeline.

The principal assumption is that the studies conducted have sampled the microbiome within a time duration of high interest. For example, [8] are interested in looking purely at the infant gut microbiome from birth to when they are ready to be discharged from the ICU. We assume that the scientific interest purely lies in the period between the starting event (sampling point) and the ending event (sampling point) and not in the actual clock/calendar time. As a result of this, our method analyses the TVAP patterns within the time duration of interest. We pass the burden of responsibility to users to use data captured within a duration of interest.

Some of the auxiliary assumptions are:

- that the TVAP is uniform between the sampling points;
- that the microbial variation patterns are independent of/minimally affected by external influences;
- that different host environments’ microbial communities may have time lags and slower or faster dynamics;
- that OTU TVAP patterns are meaningful when considered independent of the other OTUs.

However, we do not involve some assumptions usually taken with microbial community dynamics analysis, including the assumption of a particular pattern for an OTU behaviour.

### 3.4 Knowledge from precision medicine

Precision medicine is where an individual’s specialised needs are considered before prescribing the medicine. Although this seems trivial, not all individualities are considered in medicine, especially when microbial individualities are concerned. There is a growing interest in gut treatments to consider the composition and dynamics of an individual’s gut microbial ecosystem before prescribing medicine.

Furthermore, research suggests that other ailments also could be treated through the gut microbiome targeted precision medicine therapy [25]. Use of the same antibiotics and probiotics as a generalised therapy is undesirable if the patients’ gut microbial compositions are entirely different, as we can reasonably expect that the reaction to anti/probiotics would vary for different microbial communities. Hence, an analysis that exposes the generalisation level applicable to individuals is a requirement for precision medicine.

### 3.5 Intra-cluster variations / sub-clusters

In some cluster configurations, we also observe sub-clusters. This observation, we propose to indicate that even in the apparent intra-cluster conformity, some individualistic traits prevail. These subclusters, especially when visualised as abundance variation patterns, can show us minute idiosyncrasies of the microbial dynamics. We believe that we would explain these subclusters in the future with enough clinical and environmental information.

### 3.6 Gammaproteobacteria & Betaproteobacteria Clusters

We identified *Gammaproteobacteria* as a major OTU in the gut and a secondary OTU in the respiratory microbiome. *Betaproteobacteria* was identified as a major OTU in the respiratory microbiome. The behaviour of the *Proteobacteria* in the visualisation was intriguing. Summarily, they almost always showed clearly separated and tight TVAP pattern clusters. Both *Gammaproteobacteria* and *Betaproteobacteria* were often the most clearly separated in many datasets, which was consistent in the datasets we examined.

### 3.7 Separation of Clusters at Different Taxonomic Resolutions

We observe the separation of clusters and indications of individuality and conformity at all taxonomic resolutions. We propose that this indicates that our visualisation is not necessarily a good technique only at the class level but also at other taxonomic levels. We observe the preserved structure in Figure 11 across the taxonomic hierarchy of Phylum *Firmicutes*, Class *Bacilli*, Order *Lactobacillales*, Family *Streptococcaceae*, and Genus *Streptococcus*. We propose that we interpret this preservation of structure as a trait (related to microbial dynamics) that is similarly observed in closely related microbial species.

### 3.8 Heterogeneity and Complexity

We argued earlier that time-series microbial datasets are complex and heterogeneous. The CoPR visualisations confirm that fact. We observe that the underlying structures of the microbial abundances are not homogeneous even in similar host environments. They are also different across different OTUs in the same host environment. We also see connections, such as cluster agreement which appear across OTUs, and across host environments. These observations reinforce our argument of the complexity and heterogeneity of microbial abundance data.

### 3.9 Distinctions from Other Collective Pattern Recognition Approaches

One of the main pitfalls we identify is the assumption that a typical TVAP pattern exists. When a method strives to achieve that typical pattern, it results in a loss of information. However, with our visualisation, we have shown that such a common trait does not exist, and that assumption should be invalid.

Secondly, another distinction in our approach is that it prevents the loss of individuality. Like other approaches, we also identify common patterns. However, that conformist approach is not at the expense of loss of individuality. The dominant pattern is preserved in the other approaches while forcing other patterns to transform into it [7]. While we agree that using the DTW distance can be considered a transform, its use is always pairwise—hence, it does not give prominence to a single TVAP pattern.

### 3.10 Future Work

We have identified several future research directions made possible through the CoPR visualisations.

Firstly, we are interested in exploring the subclusters and the intricacies involved in their separation. In the future, with more clinical and environmental data, we believe this could be quite an intriguing research direction to take.

Secondly, we can exploit the median TVAP curves in IMPARO to obtain MINs for each cluster combination. Because each OTU has multiple clusters, there would be multiple MINs inferred. However, this is in agreement with the idea presented in IMPARO [1] that we cannot infer a single solution for MIN by examining NGS data. Hence, observations from the CoPR pipeline and observations from IMPARO directly complement each other. By identifying cluster overlaps that are statistically significant through the CoPR pipeline, we can improve the interpretation of MINs acquired through IMPARO. Isolation of overlapping clusters will help identify the behavioural/interaction patterns of OTUs in a homogeneous subset of environments, assisting in developing a global picture of OTU interactions where ambiguity is minimised. These findings may shine a light on separating environmental factors from MINs as well. We identify and this task as an exciting future research direction. Thirdly we propose it would be interesting to observe and characterise the meaningful differences in the TVAP clustering patterns of major, secondary and rare OTUs. Especially if we observe different host environments, we might be able to find whether OTUs have different temporal dynamics when they are a major OTU or not.

Fourthly, we propose to investigate a connection between the notion of dynamic microbial interaction networks and CoPR, where we consider time-windows of a lengthy dataset (such as [5]) to be a different host environment. Thus, we can apply collective pattern recognition techniques to a single abundance profile and analyse the temporal dynamics of microbial interactions through the clusters. Primarily, this could help us explore the repeated MINs discussed in [35].

Lastly, further analysis of the temporal variations of abundance profiles can be performed through a biological viewpoint. It would be of interest to examine the biological significance of the emerged patterns and the causal relationships as identified through the CoPR pipeline.

## 4 Conclusion

The TVAP patterns show that microbial community activity is heterogeneous and complex. We conclude that the behaviours of different OTUs across host environments vary and is best explored on a case by case basis. As per our discussion, there is a balance of conformity and individuality in the TVAP patterns. We propose that this behaviour can be an informative characterisation of OTU communities. We presented CoPR, a visualisation framework for collective pattern recognition for microbial data. Through unsupervised clustering of the data, our visualisation approach provided an exciting insight into the microbial communities. We believe this kind of analysis would be ideal for analysing new datasets. We also raise the question that the TVAP patterns may be connected with clinical factors. This question, however, remains to be fully answered in the future.

## 5 Methods

In this section, we first introduce the datasets we processed. Then we explain the pipeline in detail, including the techniques used and the terminology used throughout the paper.

### 5.1 Datasets

The following two datasets were used in our analysis.

#### 5.1.1 Neonatal Infant Gut Microbial Dataset

The first dataset we process is an Infant Gut Microbial dataset collected by [8]. This longitudinal dataset consists of 58 subjects, with an average of 16-time points each. Each subject is an infant in an intensive care unit. Stool samples were collected from each infant during their stay, and we have access to the abundance profiles generated through 16S rRNA sequencing and several clinical information about the infants, such as milk consumption, post-conception age, and delivery method. In our analysis of our dataset, we try to observe whether there is a connection between the clinical factors and the Temporal Variation of Abundance Profile (TVAP) patterns.

#### 5.1.2 Infant Gut and Respiratory Microbial Dataset

The second dataset we look at in this paper is another Infant Microbial dataset collected by [11]. This longitudinal abundance data set has data from 82 infants, of whom 38 are pre-term and 44 are full term. We also have data from multiple body sites. Communities from the nasal, throat and gut microbiome are analysed, contrasted with and compared to each other.

#### 5.1.3 Data Simulation

After testing the CoPR pipeline with real-life datasets, we used simulated data to verify our analysis. In this section, we will look at how I simulated the data. The dataset is created to approximate a known group with the clustering, as we identified correlations in the real-life datasets. I used a stencil-based approach, where a stencil is defined through a mathematical function. Twenty such stencils were defined. These included ten behavioural functions that defined major OTU behaviour after typical temporal behaviours examined in high abundant OTUs. A further ten reflected rare OTUs’ temporal behaviour for minor/secondary OTUs. In simulating the data, each OTU population were probabilistically assigned stencils to follow. For example, each OTU–A’s population had a probability of 0.4 to follow behaviour no. 1, 0.59 for behaviour no. 2, and 0.001 for behaviour nos. 11 to 20. TVAP were calculated according to these pre-defined behavioural stencils, to which uniform noise was added to complete the simulation.

### 5.2 Application Pipeline

In this subsection, we discuss the application pipeline we use in our work. The pipeline’s input is microbial abundance data, and the output is the visualisation. Figure 14 illustrates the different parts of this pipeline. We then discuss each of the techniques we have used in the pipeline.

**Figure 14.**
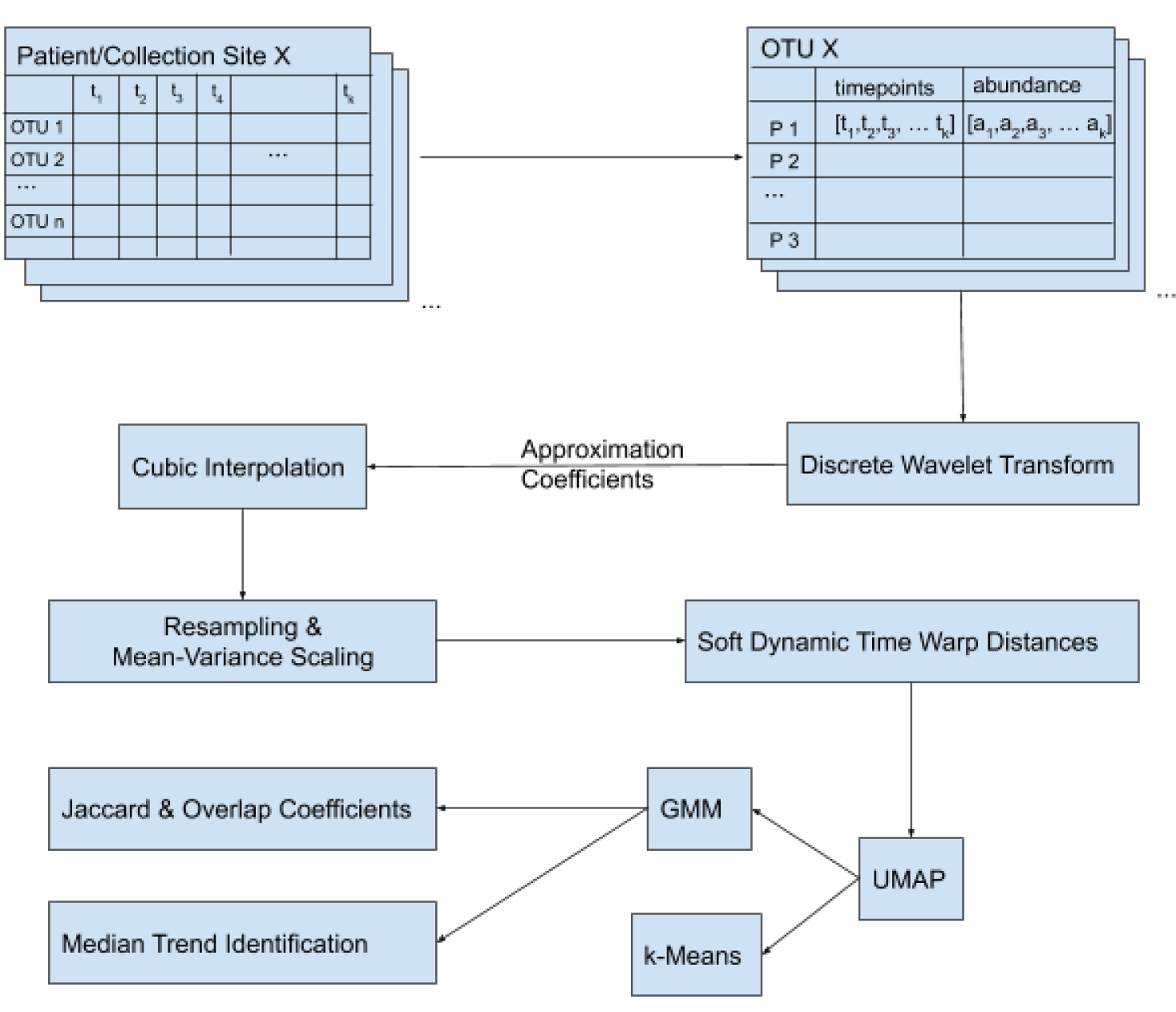
The first step is to convert the abundance data in a host-environment-wise into an OTU-wise configuration. For each host environment in the original data, we have an OTU abundance tabled against time points. We have a two-column table for each OTU in the new data where each row represents a patient/collection site. The two columns correspond to the sampling time points and the corresponding OTU abundance. The next step is to send the time-series abundance samples through a symmetrical Discrete Wavelet Transform (DWT). We then take the Approximation Coefficients and perform a cubic interpolation. After the interpolation, all the time-series are resampled to the same timeline and re-centred with respect to mean and variance. After these pre-processing steps, pair-wise soft dynamic time warp distances are calculated. These distances are used in place of the dimensions for the UMAP dimension reduction. The dimension reduced data is clustered with a Gaussian Mixed Model clustering, where the optimal cluster number has been identified through a silhouette score calculation. We also provide a k-Means clustering visualisation in the interest of comparison. As auxiliary visualisations, we provide the median trend for each cluster and selection and Jaccard and Overlap Coefficients of clusters across the major OTUs.

The first step is to convert the abundance data in a host-environment-wise configuration into an OTU-wise configuration. For each host environment in the original data, we have an OTU abundance tabled against time points. We have a two-column table for each OTU in the new data where each row represents a patient/collection site. The two columns correspond to the sampling time points and the corresponding OTU abundance. The next step is to send the time-series abundance samples through a symmetrical Discrete Wavelet Transform (DWT). We then take the Approximation Coefficients and perform a cubic interpolation. After the interpolation, all the time-series are resampled to the same timeline and re-centred with respect to mean and variance. After these pre-processing steps, pair-wise soft dynamic time warp distances are calculated. These distances are used in place of the dimensions for the UMAP dimension reduction. The dimension reduced data is clustered with a Gaussian Mixed Model clustering, where the optimal cluster number has been identified through a silhouette score calculation. We also provide a k-Means clustering visualisation in the interest of comparison. As auxiliary visualisations, we provide the median trend for each cluster and selection and Jaccard and Overlap Coefficients of clusters across the major OTUs.

#### 5.2.1 Discrete Wavelet Transform

Discrete Wavelet Transform(DWT) is the first pre-processing step. DWTs conserve both frequency and location (time in temporal data) information; hence it is more suitable for our task than a Fourier Transform. We use a symmetrical kernel filter in our DWT step. A wavelet transform can act as a high-pass and low-pass filter. We use this quality to remove the noise and small fluctuations, to obtain the general trend we are interested in. Hence after the DWT, we discard the detailed coefficients and keep the approximation coefficients to represent the time-series data. The DWT also increases the frequency resolution of the data. We used the PyWavelets implementation of DWT in our application pipeline [36].

#### 5.2.2 Cubic Interpolation

As the second pre-processing step, we use a 1D cubic interpolation. The interpolation aims to fill in the gaps between the sampling time points, as we require a continuous abundance variation pattern for comparison across host environments. Each host environment dataset is interpolated from its first time point to the last, with no extrapolation. Most other methods tend to use a spline interpolation; however, as our effort focuses on identifying an overall pattern, we consider a 1D interpolation to be more suitable in contrast with temporally localised patterns. We are assuming that the abundance pattern variation in between the sampling points is uniform.

#### 5.2.3 Mean-Variance Scaling

As a third pre-processing step, we scale each interpolated time series to be centred around the mean, with a variance of 1 (*µ* = 0, *σ*= 1). We aim to isolate each OTU community’s abundance pattern from the rest of the host environment by doing this preprocessing step. As we are merely interested in the temporal variation pattern of each OTU across multiple host environments, this allows direct comparison. The effect of this step is especially prominent in host environments, where there exist two dominant OTUs. This pre-processing step will hinder any quantification of microbial interactions. Hence, it is crucial to reiterate that we are not interested in the microbial interactions in this visualisation pipe-line, but rather the abundance variation pattern is our interest.

#### 5.2.4 Resampling

The final step in our pre-processing approach is resampling. It is done to align the timelines of different host environments. We acknowledge that there are solid arguments for and against resampling the timelines across different datasets. The resampling will shift the timelines and change the temporal scale, which results in losing temporal information. However, when the sampling is done in a meaningful time scale, the resampling can help find a better overlap across samples. Hence the choice of resampling aligns with our choice of using dynamic time warp distance as well.

#### 5.2.5 Dynamic Time Warp Distance

Dynamic Time Warp (DTW) Distance has been used in many time-series clustering-based methods in the literature. DTW is best explained as the distance between two-time series at their best temporal alignment. We seek a temporal alignment as different host environments could have delayed or temporally inconsistent behaviour, which can be identified to be based on similar variation patterns. By using the DTW distance, we can cluster similar variation patterns, despite temporal inconsistencies. We use the tslearn [37] implementation of the DTW distance in our application pipeline.

#### 5.2.6 UMAP

Uniform Manifold Approximation and Projection (UMAP) [38] is a manifold learning technique for dimension reduction. It is considered to have high visualisation quality and preserve more global structure than other dimension reduction techniques such as t-SNE [39]. We use UMAP to reduce the temporal dimensions and visualise each OTU’s TVAP as a point in a 2-D plane. The neighbourhoods are determined by the DTW distance between each pair of datasets. The UMAP visualisation gives us an idea about the similarities and differences between time series data sets. We can observe that points that are clustered together correspond to similar TVAP patterns.

#### 5.2.7 GMM Clustering

We cluster the data-points in the reduced dimension using a Gaussian Mixed Model (GMM) clustering. The number of clusters was determined by calculating the Silhouette Score for each cluster configuration. While we also consider k-Means clustering, GMM clustering results in superior identification of clusters. Because GMM considers the variation and the mean for its clustering, GMM more accurately identifies cluster membership of adjacent clusters of different sizes.

#### 5.2.8 Silhouette Score

Silhouette score is a measure of how similar an object is to its own cluster and how different it is compared to the objects in other clusters (cohesion vs separation). This is a useful visual aid of the clustering performance [40]. The silhouette score graph we presented is the silhouette score as a function of the number of clusters. By examining the silhouette score graphs, we can understand how distinct the separation is and how similar the cohesion is at different cluster numbers. The higher the silhouette score is, the better the cohesion and separation. We calculate the silhouette score for the same number of clusters multiple times and take the average. This calculation also provides us with the standard deviation (shown with the error bar) for the silhouette scores. A narrower error bar means that the clustering is consistent at that number. A high silhouette score with a narrower error bar is the ideal cluster configuration we are looking for.

#### 5.2.9 Overlap Coefficient & Jaccard Index

We use both the Overlap Coefficient (Szymkiewicz-Simpson Coefficient) (Equation 1) and the Jaccard Index (Equation 2) to examine the corresponding behaviour among the clusters. While the Jaccard Index indicates a bi-directional correspondence amongst two clusters, using the Overlap Coefficient allows us to identify uni-directional correspondence.

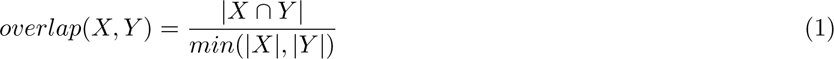

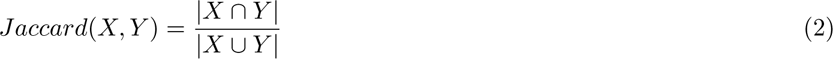

#### 5.2.10 Bokeh Visualisation Engine

Finally, we use the Bokeh Visualisation Engine [41] to provide interactivity to the visualisations, allowing a thorough manual exploration of the clusters and associated TVAP patterns.

### 5.3 Terminology

In this subsection, we will introduce some terminology which we have used throughout the paper.

#### 5.3.1 TVAP

Temporal Variation of abundance Profile. When we look at the abundance of a certain OTU as a function of time, we get the abundance profile’s temporal variation. We have obtained a continuous graph by interpolating between the sampling points. This function, when visualised, will show certain tendencies. These tendencies are what we would regard to be identifiable patterns specific to each OTU community. As an example, we may notice rising abundance patterns, dropping abundance patterns, sudden peaks, etc. We argue in this paper that we can use these patterns to characterise an OTU community.

#### 5.3.2 OTU community

For this paper’s scope, we characterise an OTU community as the organisms of a specific Operational Taxonomic Unit, which inhabit a particular host environment. To illustrate, we consider *Gammaproteobacteria* in a specific infant’s gut microbiome as an OTU community. There could be several *Gammaproteobacteria* communities in the same infant, such as the gut *Gammaproteobacteria* community and the respiratory *Gammaproteobacteria* community. For the purpose of this paper, we consider them to be two separate and independent OTU communities.

#### 5.3.3 Major OTU

In order to efficiently compute the CoPR analysis pipeline, we choose a subset of OTUs. In most analyses, the interest would be on the most abundant OTUs. Hence, consider the *n* highest abundant OTUs of each host environment in terms of average abundance. For this work’s scope, let us define major OTUs as the intersection of the sets of *n* highly abundant OTUs in all the parallel host environments. Likewise, let us define secondary OTUs as the OTUs in the top 2*n*, excluding the major OTUs.

The parameter *n* can be set according to the need of the analysis.

## Availability of data and materials

Real-life data used in this study is publicly available at MG-RAST:4457768.3-4459735.3. The code of CoPR is available at https://bitbucket.org/rajith/copr/, and is released publicly under MIT license. All simulated data sets, and python notebooks are also available in the repository.

